# Feasibility of intranasal delivery of thin-film freeze-dried, mucoadhesive AS01_B_-adjuvanted vaccine powders

**DOI:** 10.1101/2022.11.01.514748

**Authors:** Yu-Sheng Yu, Khaled AboulFotouh, Gerallt Williams, Julie Suman, Chris Cano, Zachary N. Warnken, Robert O. Williams, Zhengrong Cui

## Abstract

Intranasal vaccination by directly applying a vaccine dry powder is appealing. However, a method that can be used to transform a vaccine from a liquid to a dry powder and a device that can be used to administer the powder to the desired region(s) of the nasal cavity are critical for a successful intranasal vaccination. In the present study, using a model vaccine that contains the liposomal AS01_B_ as an adjuvant and ovalbumin (OVA) as a model antigen, it was shown that thin-film freeze-drying can be applied to convert the liquid vaccine containing sucrose at a sucrose to lipid ratio of 15:1 (w/w), in the presence or absence of carboxymethyl cellulose sodium salt (CMC) as a mucoadhesive agent, into dry powders. Ultimately, the thin-film freeze-dried AS01_B_/OVA vaccine powder containing 1.9% w/w of CMC (i.e., TFF AS01_B_/OVA/CMC_1.9%_ powder) was selected for additional evaluation because the TFF AS01_B_/OVA/CMC_1.9%_ powder was mucoadhesive and maintained the integrity of the antigen and the physical properties of the vaccine. Compared to the TFF AS01_B_/OVA powder that did not contain CMC, the TFF AS01_B_/OVA/CMC_1.9%_ powder had a lower moisture content and a higher glass transition temperature and was more porous. In addition, the TFF AS01_B_/OVA/CMC_1.9%_ thin films were relatively thicker than the TFF AS01_B_/OVA thin films without CMC. When sprayed with the Unit Dose System Powder (UDSP) nasal device, the TFF AS01_B_/OVA powder and the TFF AS01_B_/OVA/CMC_1.9%_ powder generated similar particle size distribution curves, spray patterns, and plume geometries. Importantly, after the TFF AS01_B_/OVA/CMC_1.9%_ powder was sprayed with the UDSP nasal device, the integrity of the OVA antigen and the AS01_B_ liposomal adjuvant did not change. Finally, a Taguchi L8 orthogonal array was applied to identify the optimal parameters for using the UDSP device to deliver the TFF AS01_B_/OVA/CMC_1.9%_ vaccine powder to the middle and lower turbinate and the nasopharynx regions in both adult and child nasal casts. Results from this study showed that it is feasible to apply the TFF technology to transform a nasal vaccine candidate from liquid to a dry powder and then use the UDSP nasal device to deliver the TFF vaccine powder to the desired regions in the nasal cavity for intranasal vaccination.

## 1. Introduction

Intranasal vaccination is an attractive, non-invasive route of vaccine administration. Intranasal vaccine can induce specific immune responses not only systemically, but also in the mucosal secretions of the respiratory tract (Birkhoff et al., 2009), which is advantageous as many pathogens infect their hosts through the respiratory tract (Chavda et al., 2021). In fact, several nasal vaccines have been approved for human use around the world, including the FluMist Quadrivalent in the United States (US) (Suryadevara and Domachowske, 2014), Fluenz Tetra in the European Union (EU) (Gasparini et al., 2021), NASOVAC-S in India (Ortiz et al., 2015), a freeze-dried nasal live influenza vaccine (Ganwu^®^) by the Changchun BCHT Biotechnology in China, and recently a nasal Coronavirus disease 2019 (COVID-19) vaccine by Bharat Biotech in India. Those nasal vaccines are live (attenuated) influenza virus-based or adenovirus-based and are presented as a liquid suspension or a freeze-dried powder for reconstitution. The liquid vaccine suspension is then administered intranasally using a nasal sprayer. Intranasal administration of vaccine directly as a dry powder has advantages, including ease of storage and distribution (Flood et al., 2016), extended residence time in the nasal cavity, and a higher dose of vaccine that can be administered. Unfortunately, vaccine dry powders with the proper aerosol properties for deposition in the desired region(s) in the nasal cavity and a method to prepare the dry powders remain needed.

Thin-film freeze-drying is a bottom-up dry powder engineering technology. It involves the ultra-rapid thin-film freezing (TFF) of a liquid (e.g., solution, suspension, or emulsion) on a cryogenically cooled solid surface (Engstrom et al., 2008; Overhoff et al., 2009). The liquid is dropped as droplets (e.g., 2 mm in diameter) from about 1 cm to 10 cm above the cryogenically cooled solid surface. Upon impact onto the cooled surface, the droplet rapidly spreads into a thin film, which is then frozen into a thin film. Solvent such as water in the frozen thin films is removed by lyophilization. Dry powder engineering using the TFF technology is advantageous over other similar dry powder engineering technologies such as conventional shelf freeze-drying, spray drying, and spray freeze-drying in that it avoids or minimizes shear stress and heat stress, while generating powders that are generally highly porous, brittle (Hufnagel et al., 2022), and having large specific surface area (Wang et al., 2021), making them potentially feasible for directly intranasal administration. Conversely, the TFF process is associated with a relative larger liquid-solid interface between the liquid droplets and the cryogenically cooled solid surface. Previously, it has been shown that the TFF technology can be applied to produce dry powders of various vaccines, including vaccines adjuvanted with aluminum salts (Alzhrani et al., 2021; Li et al., 2015; Thakkar et al., 2018), (nano)emulsions such as MF59 or AddaVax (AboulFotouh et al., 2022a), and liposomes such as AS01_B_ (AboulFotouh et al., 2022b).

Herein, a model vaccine comprised of ovalbumin (OVA) as a model antigen and the liposomal AS01_B_ as an adjuvant was used to test the feasibility of using the TFF technology to prepare dry powder vaccines for direct intranasal administration. All currently approved human nasal vaccines are virus-based and do not contain any adjuvant. The AS01_B_ was chosen in this study because a recent report from our group showed that AS01_B_-containing vaccines can be converted to dry powders by TFF (AboulFotouh et al., 2022b). Results from a recent study have demonstrated the safety and efficacy of AS01_B_ as a potential nasal vaccine adjuvant in a mouse model (Sato-Kaneko et al., 2022). Moreover, one can find in the literature intranasal immunization studies wherein the vaccines contain either monophosphoryl lipid A (MPL) or QS21 as the adjuvant (Baldridge et al., 2000; Sasaki et al., 1998a; Sasaki et al., 1998b), and the liposomal AS01_B_ contains both MPL and QS21.

To increase the residence time of the vaccine in the nasal cavity, various mucoadhesive agents have been included in the vaccine formulation, and their effect on the vaccine was studied. The mucoadhesive agents studied include chitosan, sodium alginate, gelatin, and sodium carboxymethylcellulose (CMC), each with its own unique mechanism(s) of interaction with the mucosa (Dekina et al., 2016; Grabovac et al., 2005; Kesavan et al., 2010; Sogias et al., 2008). A challenge is that the mucoadhesive agent can interact with the AS01_B_/OVA vaccine and change the structure of the AS01_B_, the antigen, and/or the vaccine. Therefore, the effect of the mucoadhesive agents on the AS01_B_/OVA vaccine before and after being subjected to thin-film freeze-drying with sucrose as an excipient was studied. One of the mucoadhesive agents at a concentration that showed no detectable effect on the physical properties of the AS01_B_/OVA vaccine was chosen for further characterization. Aptar’s unit dose system powder (UDSP) nasal device was used to characterize the particle size distribution, spray pattern, and plume geometry of selected AS01_B_/OVA vaccine powders upon actuation. Finally, the deposition patterns of a selected AS01_B_/OVA vaccine powder following actuation using the UDSP nasal device into 3D printed nasal casts based on the CT-scan images of the noses of an adult and a child were evaluated to predict the feasibility of using the UDSP nasal device to deliver thin-film freeze-dried vaccine powders (TFF vaccine powders) directly into the posterior nasal cavity and the nasopharynx regions of human nasal cavity (Warnken et al., 2018). Unlike in infants, in children older than two years and in adults, the Waldeyer’s ring in the naso-oropharynx region is the key lymphoid tissue in the nasal cavity (Davis, 2001; Debertin et al., 2003).

Therefore, for an intranasal vaccine to induce specific immune responses, the vaccine needs to be delivered either directly to the nasopharynx region or to the posterior nasal cavity and then move by means of ciliary clearance to the naso-oropharynx region following the mucous blanket posterior movement. Within the posterior nasal cavity, delivery of the vaccine to the upper turbinate region, where the olfactory bulb resides, should be avoided or minimized to reduce the access of the vaccine to the brain via the olfactory nerves (Cai et al., 2022; Xu et al., 2021b).

## 2. Materials and Methods

### 2.1. Materials

Albumin from chicken egg white (ovalbumin, OVA), chitosan medium molecular weight, sodium alginate, gelatin, CMC, MPL from *Salmonella enterica* serotype minnesota Re 595 (Re mutant), porcine mucin type III, and 2-mercaptoethanol were from Sigma-Aldrich (St. Louis, MO). QS-21 was from Dessert King International (San Diego, CA). Sucrose was from Millipore (Billerica, MA). Anhydrous ethanol (EtOH) was from Decon Labs (King of Prussia, PA). Cholesterol was from MP Biomedicals (Santa Ana, CA). The 1,2-dioleoyl-sn-glycero-3-phosphocholine (DOPC) was from Avanti Polar Lipids, Inc. (Alabaster, AL). Fluorescein isothiocyanate isomer I (FITC) was from Acros Organics (Geel, Belgium). Dulbecco’s phosphate-buffered saline (DPBS, pH 7.0-7.3, 9.5 mM) was from Gibco (Grand Island, NY). Laemmli sample buffer 4× and Coomassie G-250 were from Bio-Rad (Hercules, CA). Blue prestained protein standard was from New England Biolabs (Ipswich, MA). Artificial nasal mucus was from Biochemazone (Leduc, Alberta, Canada). HYDRANAL™ - Coulomat AG was from Honeywell (Charlotte, NC). The UDSP nasal device was kindly donated by AptarGroup, Inc. (Crystal Lake, IL). All chemicals were used as received without further purification.

### 2.2. Preparation of the AS01_B_/OVA model vaccine

To prepare the AS01_B_/OVA model vaccine, the liposomal adjuvant was prepared first as described before (AboulFotouh et al., 2022b). Briefly, 4.0 mg of DOPC, 1.0 mg of cholesterol, and 0.2 mg of MPL were dissolved with 1.25 mL of EtOH in a 20 mL glass vial. A lipid membrane was formed in the vial by evaporating the ethanol under gentle nitrogen flow. Next, 0.5 mL of DPBS was added to the vial to rehydrate the lipids, and the mixture was stirred overnight at room temperature. The final concentration of DOPC in the liposome suspension was 8 mg/mL. The liposome suspension was sonicated with a probe sonicator (Qsonica Q700) until the particle size reached about 100 nm based on dynamic light scattering (DLS). The AS01_B_/OVA model vaccine was prepared by adding 50 μg of QS-21, 50 μg of OVA, and 19.5 mg of sucrose to 125 μL of the liposome suspension, and the final volume was adjusted to 500 μL with DPBS.

### 2.3. Preparation of AS01_B_/OVA model vaccines with different mucoadhesive agents

To prepare the AS01_B_/OVA model vaccine with different concentrations of chitosan, sodium alginate, gelatin, or CMC, 50 μg of QS-21, 50 μg of OVA, and 19.5 mg of sucrose were first added to 125 μL of liposome suspension. The stock solution of chitosan was prepared by dissolving chitosan in a 0.1 M acetic acid aqueous solution (2% w/v). The stock solutions of sodium alginate, gelatin, and CMC were prepared by dissolving them in DPBS (2% w/v). Different volumes of the solutions containing mucoadhesive agents were then added to the AS01_B_/OVA vaccine to achieve a final concentration of 0.1%, 0.2%, 0.4%, 1%, w/v, corresponding to 1.9%, 3.7%, 7.2%, and 16.3% (w/w) of the mucoadhesive agents to the total weight of all components in the vaccine formulation except water. The final volume of the AS01_B_/OVA vaccines was then adjusted to 500 μL with DPBS.

### 2.4. Thin-film freeze-drying of the AS01_B_/OVA model vaccines

The as-prepared AS01_B_/OVA vaccines, with or without mucoadhesive agents, were converted into dry powders by the TFF process. First, the liquid vaccines were frozen into small thin films following the reported single-vial TFF process (Xu et al., 2021a). Briefly, a 20 mL glass vial was placed on dry ice until the temperature reached equilibrium. The liquid vaccine was added dropwise with a BD 1 mL syringe with a 21G needle to the bottom of the vial to form frozen thin films. The frozen films were then freeze-dried in a lyophilizer (SP VirTis AdVantage Pro). The lyophilization cycle was -40°C for 20 h, gradually ramping from -40°C to 25°C in 20 h, and then 25°C for 20 h. The chamber pressure was maintained at 80 mTorr. After lyophilization, the vials were backfilled with nitrogen gas, capped, crimped, and stored at room temperature.

### 2.5. Determination of particle size and zeta potential

A Malvern Nano ZS (Westborough, MA) was used to measure the particle size and zeta potential of the liposomes and the vaccines. The vaccine powder was reconstituted with deionized water (DI water) at a volume identical to the volume of the liquid vaccine before it was subjected to TFF. Samples were diluted 50 times with DPBS. The final pH value of all the samples was approximately 7.2 to 7.4.

### 2.6. Evaluation of the integrity of the OVA and the liposome

Sodium dodecyl sulfate–polyacrylamide gel electrophoresis (SDS-PAGE) was applied to evaluate the integrity of OVA in AS01_B_/OVA vaccines before and after they were subjected to thin-film freeze-drying, as well as before and after the vaccine powders were sprayed with an Aptar UDSP nasal device. Vaccine powders were reconstituted with water. To spray a powder, the powder was filled into a UDSP nasal device following the manufacturer’s instruction and sprayed into a glass pipette rubber head. The powder was then reconstituted from within the rubber head with water. For SDS-PAGE analysis, the sample was mixed with Laemmli sample buffer 4× containing 10% 2-mercaptoethanol, boiled for 10 min at 100°C, and then loaded into a well of a 4–20% precast polyacrylamide gel from Bio-Rad (Hercules, CA) (30 μL per well). The blue prestained protein standard was also loaded to the gel. The electrophoresis was performed at 90 V for 90 min. The SDS-PAGE gel was stained with Coomassie G-250, and the gel image was captured with a camera.

Scanning transmission electron microscope (STEM) was utilized to examine the integrity of the liposomes after the vaccine powder was sprayed with a UDSP nasal device. The vaccine powder was reconstituted and diluted 5 times with water and then transferred to a glow discharge-treated carbon coated copper grid from Electron Microscopy Sciences (Hatfield, PA). The sample was stained with a 2% uranyl acetate solution for 2 min and then air dried. Finally, the STEM images were captured with a Hitachi S5500 (Tokyo, Japan) available in the Texas Materials Institute at UT Austin.

### 2.7. In vitro mucoadhesion test

The *in vitro* mucoadhesion test was done following a previously reported procedure with modifications (Trenkel and Scherließ, 2021). To simulate the human nasal mucosal tissue, 10 cm Petri dishes were coated with two different solutions: 1.5% (w/v) agar in DPBS (pH = 6) or 1.5% (w/v) agar plus 2% (w/v) porcine mucin in DPBS (pH = 6). The coating layer was solidified by incubating the Petri dishes for 2 h at room temperature followed by 30 min at 4°C. After solidification, the coated dishes were incubated at 32°C in a Fisher Scientific incubator-shaker (Hampton, NH) until equilibrium (shaking not applied), and 3-5 mg of the vaccine films were gently placed onto the gel. The dishes were then turned vertically and incubated for 2 h at 32°C. The maximum downward movement of the vaccine powders at the time points of 10 min, 20 min, 30 min, 1 h, and 2 h were recorded.

### 2.8. Characterization of the TFF vaccine powders

XRD was done using a Rigaku R-Axis Spider (Tokyo, Japan) available at UT Austin. Vaccine powder was mounted in mineral oil and loaded onto the sample loops. The powder was irradiated with Cu Kα radiation, and the diffraction pattern was captured with a 2-D detector. The 2-D pattern was then converted to 1-D pattern. The crystal structures of the sodium chloride (NaCl) and sucrose were obtained from the Crystallography Open Database (COD) (http://www.crystallography.net), and the reference patterns were generated with a Cambridge Crystallographic Data Center (CCDC) Mercury software. Also, the crystallite sizes (τ) of the samples were calculated using the Scherrer equation:

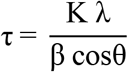

Where K is the shape factor 0.9, λ is the Cu Kα radiation wavelength, β is the full width at half maximum (FWHM) of the peaks, and θ is the Bragg angle.

To obtain the scanning electron microscope (SEM) images of a sample, a film of the TFF AS01_B_/OVA or the TFF AS01_B_/OVA with 1.9% CMC was placed onto the conductive carbon tape. The sample was then sputtered with a layer of Au/Pd (60:40) (40 mA, 1 min). SEM images were taken with a Hitachi S5500 with an acceleration voltage of 20 kV.

The surface profiles of the thin films were measured with Keyence VK-X1100 profilometer (Osaka, Japan) available at UT Austin. Before the measurement, a vaccine thin film was placed onto the unpolished side of the silicon wafer with the top up. The 3D reconstructed images of the film were captured with a 2.5 × objective lens while the ring light source was utilized. The thickness of the film was calculated with the Keyence Multi File Analyzer software.

The residue water content in vaccine powder was determined by Karl Fischer titration using a Mettler Toledo C20 coulometer (Columbus, OH). Briefly, 2.5 mL of Coulomat AG solution was drawn from the solution tank and injected into the vial containing the powder with a gas-tight syringe. The solution was mixed well with the powder, and 2.0 mL of the mixture was then injected back to the solution tank. The amount of water in the samples was then determined by the instrument, and the water content (%) in the samples was calculated with the following equation:

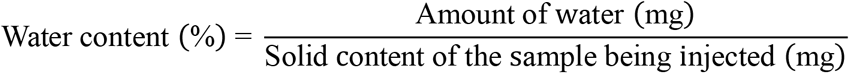

Temperature-modulated differential scanning calorimetry (mDSC) was performed using a TA Instruments Q20 differential scanning calorimeter (New Castle, DE) available at UT Austin. Briefly, 3-5 mg of the powder was loaded to the Tzero hermetic pan from TA Instruments. The scanning range was from -20 to 250°C. The temperature was modulated ±1.0°C every 60 s. The data were analyzed with the TA Instruments TRIOS software.

### 2.9. Plume geometry, spray pattern, and the particle size distribution of the aerosolized powder

The spray pattern and the plume geometry of the TFF vaccine powders were captured with an in-house built laser sheet system (Warnken et al., 2018). To study the plume geometry of a vaccine powder, the powder was loaded into a UDSP nasal device and sprayed upward.

Images were captured with a camera at a rate of 500 frames per second. To identify the boundary of the plume, the Fiji software (https://imagej.net/software/fiji) was used to create the z-stacking images from different time points to measure the plume angle. For the spray pattern, the vaccine powder was loaded into a UDSP nasal device and sprayed upward, and the laser sheet was located 6 cm above the tip. The images were captured at an angle of 45° and a frame rate of 500 frames per second. The Fiji software was then used to create the z-stacking images, and the perspective correction of the images was performed with MATLAB R2021b software. The ovality and the area were then measured with the Fiji software.

To measure the particle size distribution of the sprayed vaccine powder, the powder was filled into a UDSP nasal device and sprayed to measure the particle size with Malvern Spraytec. During the experiment, the device was fixed with a clamp, and the angle was adjusted to 30° above to the horizontal plane. The distance between the tip and the laser spot was 6 cm, while the device to lens distance was 9 cm. The device was actuated manually, and the laser diffraction signal was recorded at a frame rate of 2500 Hz.

SEM was also utilized to examine the particles after the vaccine powder was sprayed with a UDSP nasal device. The SEM specimen mount was first covered with a non-porous carbon tape. The vaccine powder was then sprayed onto the carbon tape at a distance of 6 cm. The sample was then sputtered with Au/Pd (60:40) (40 mA, 1 min), and the images were captured with an acceleration voltage of 20 kV.

### 2.10. Deposition pattern of the TFF vaccine powder

For this experiment, FITC-labeled OVA (i.e., OVA-FITC) was used to prepare the AS01_B_/OVA-FITC vaccine with 1.9% CMC. The detailed preparation procedure of the OVA-FTIC is in Supplementary Material S1.

The two 3D-printed nasal casts based on the CT-scan images of a male of 48 years of age and a female of 7 years of age were used to perform the deposition test (Warnken et al., 2018). The inner surface of the nasal cast was coated with an artificial nasal mucus, and the excess fluid was purged with air. The TFF AS01_B_/OVA/CMC_1.9%_ vaccine powder (OVA labeled with FITC) was loaded into a UDSP nasal device and sprayed into the left nostril of the nasal cast. The coronal angle and sagittal angle were 0° or 20° and 45° or 60°, respectively. The definitions of the coronal angle and the sagittal angle are shown in **Fig. sup1**. The insertion depth was 0.5 cm for the 48-year-old male nasal cast and 0.4 cm for the 7-year-old female nasal cast. To determine the effect of breathing on the deposition pattern, the nasal cast was connected to a vacuum pump to evaluate the effect of flow rate on the deposition pattern. The flow rate was set at 0 or 10 litter per min (LPM) for the 48-year-old male nasal cast and 0 or 5 LPM for the 7-year-old nasal cast. For three parameters (i.e., coronal angle, sagittal angle, and flow rate), each with 2 levels, the Taguchi L8 orthogonal array (**Table 1**) was used to find their optimal combination.

**Table 1.**
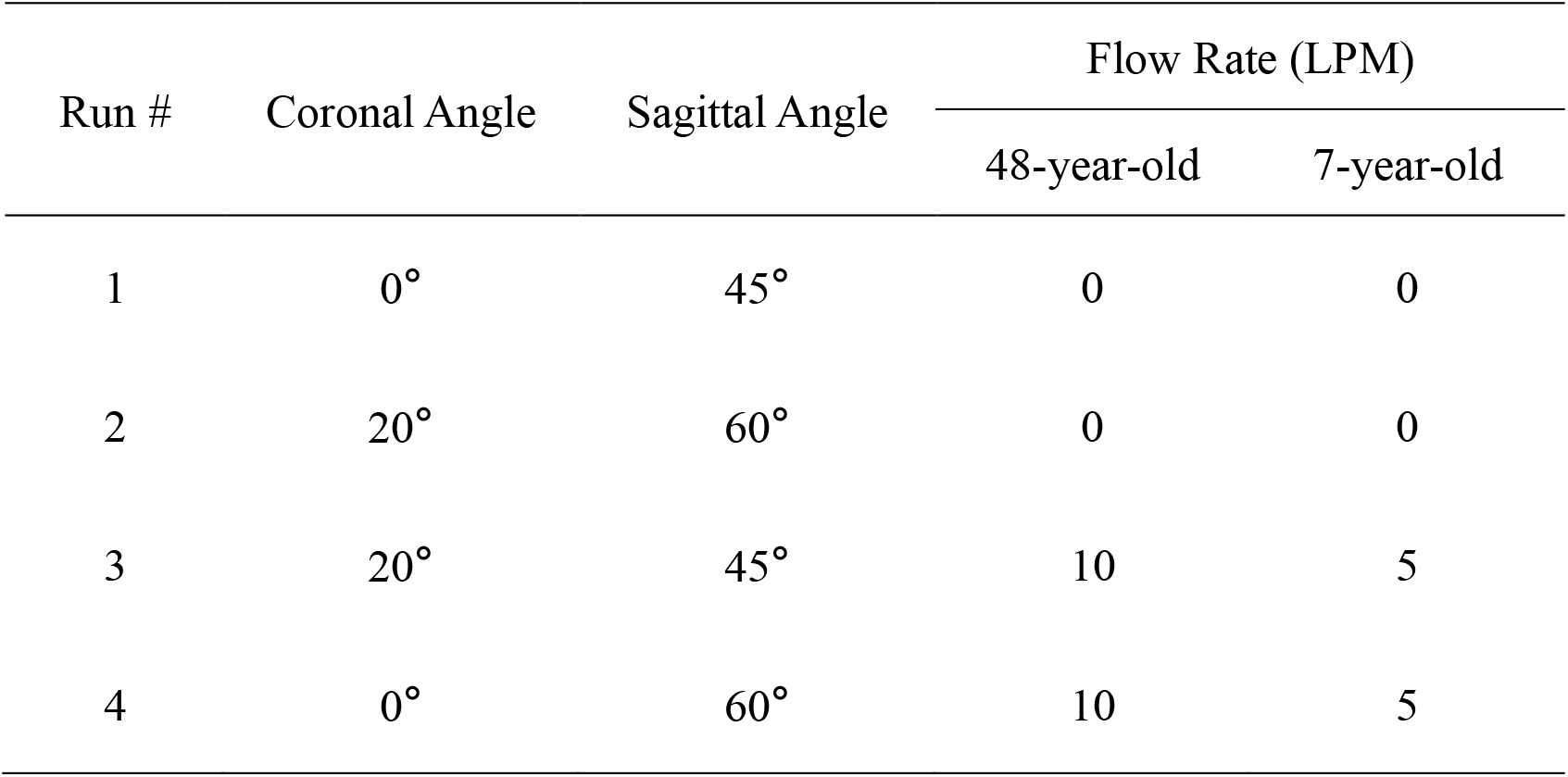
The Taguchi L8 orthogonal array for studying the deposition patterns of the TFF vaccine powder in two different nasal casts

| Run # | Coronal Angle | Sagittal Angle | Flow Rate (LPM) |  |
| --- | --- | --- | --- | --- |
|  |  |  | 48-year-old | 7-year-old |
| 1 | 0° | 45° | 0 | 0 |
| 2 | 20° | 60° | 0 | 0 |
| 3 | 20° | 45° | 10 | 5 |
| 4 | 0° | 60° | 10 | 5 |

After the powder was aerosolized, the nasal cast was disassembled into the following parts: (1) vestibule, (2) upper turbinate, (3) middle turbinate, (4) lower turbinate, (5) nasopharynx, and (6) filter. Each part was rinsed with 5 mL of water. The fluorescence intensities of the solutions were measured with a BioTek Microplate Reader (Winooski, VT). The amount of powder deposited in each part was calculated based on a calibration curve constructed with FITC-labeled vaccine powder dissolved in water.

The delivery efficiency of the powder using the UDSP nasal device was calculated based on the following equation:

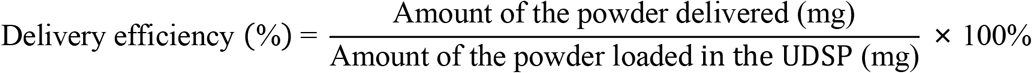

The recovery percentage of the vaccine powder in the nasal cast was defined as:

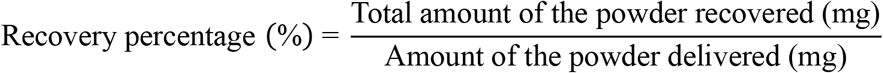

Where the total amount of the powder recovered was based on the fluorescence intensity measurement.

The deposition efficiency of the vaccine powder in a specific part or region of the nasal cast was defined as:

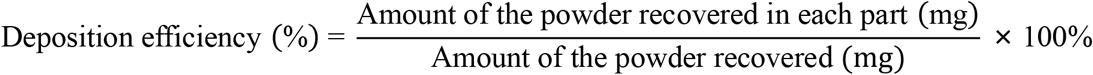

Finally, the total amount of the powder recovered in the desirable region (i.e., middle turbinate, lower turbinate, and nasopharynx region) was defined as the regional deposition efficiency (RDP).

To identify the optimized conditions for the RDP, the signal to noise (S/N) ratio was calculated for each parameter at different levels based on the “larger the better” characteristic of the Taguchi method (Ghani et al., 2004).

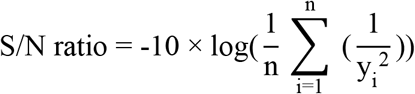

Where n is the number of observations for each parameter, and y is the average of the observation (i.e., RPD).

### 2.11 Statistical analysis

Statistical analysis was performed with Microsoft Office Excel or OriginLab OriginPro 2022b. Data are presented as mean ± standard deviation (SD), and the number of the replication (n) was indicated in the figure caption.

## 3. Results and Discussions

### 3.1. Preparation and thin-film freeze-drying of the AS01_B_/OVA model vaccine

AS01_B_ is a liposomal adjuvant comprised of DOPC, cholesterol, MPL, and QS21. AS01_B_ is prepared by adding QS21 in preprepared liposomes comprised of DOPC, cholesterol, and MPL (1:0.25:0.05, w/w) (AboulFotouh et al., 2022b). The DOPC/cholesterol/MPL liposomes prepared had a hydrodynamic diameter of 101.3 ± 5.3 nm (n = 7). The AS01_B_/OVA vaccine was then prepared by mixing the liposome suspension with a solution of OVA and a solution of QS21. Data from a previous study from our group showed that sucrose at a sucrose to lipid ratio of 15:1 (w/w) protected the AS01_B_/OVA vaccine during thin-film freeze-drying (AboulFotouh et al., 2022b). **Fig. 1** shows the particle size distribution curves of the AS01_B_/OVA vaccine before and after it was subjected to thin-film freeze-drying and reconstitution. As expected, there was not any significant change in the particle size distribution after the vaccine was subjected to thin-film freeze-drying.

**Fig. 1.**
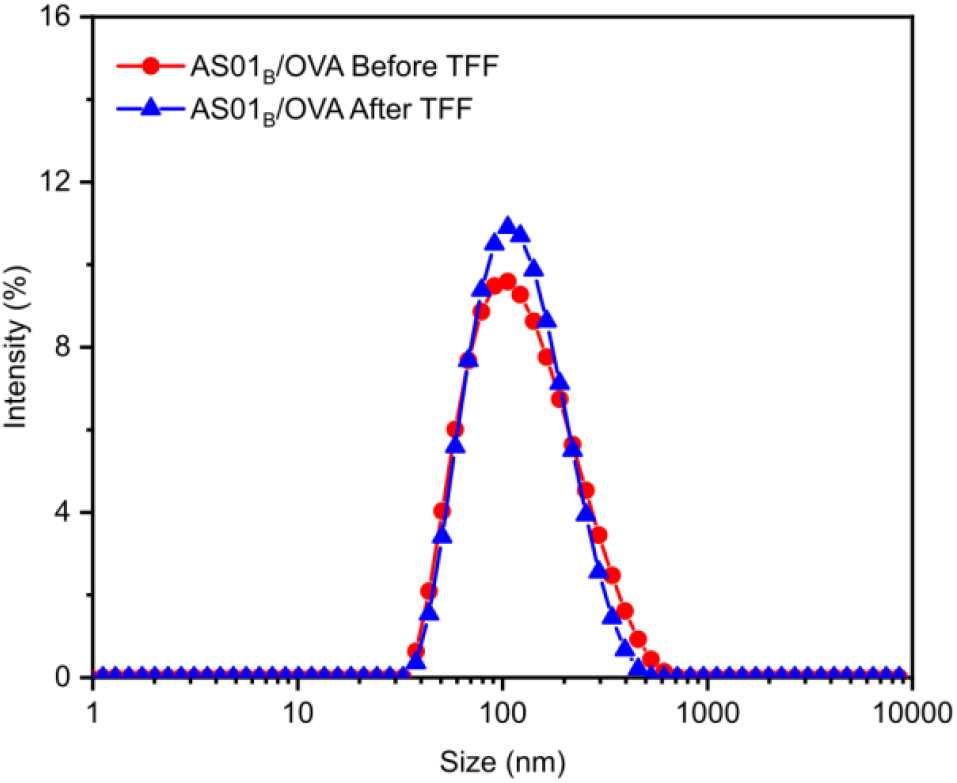
Representative particle size distribution curves of the AS01_B_/OVA vaccine before and after it was subjected to thin-film freeze-drying and reconstitution.

### 3.2. Identification of an AS01_B_/OVA vaccine formulation with a mucoadhesive agent

A mucosal adhesive agent is needed in the AS01_B_/OVA vaccine to increase its residence time in the nasal cavity. Chitosan, sodium alginate, gelatin, or CMC was added into the AS01_B_/OVA vaccine as a mucoadhesive agent to reach a final concentration of 0, 1.9%, 3.7, 7.2, or 16.3% by weight (vs. total weight of the vaccine except water), and the particle size and zeta potential values of the formulations before and after they were subjected to thin-film freeze-drying and reconstitution were determined. As shown in **Fig. sup2A**, adding sodium alginate, gelatin, and CMC at 7.2% (w/w) or below did not affect the particle size of AS01_B_/OVA vaccine, while chitosan at all concentrations tested caused a significant increase in the particle size, which might be due to the relatively small zeta potential value of the AS01_B_/OVA with chitosan (**Fig. sup2B**). After thin-film freeze-drying and reconstitution, only the AS01_B_/OVA vaccine formulations with gelatin or CMC at the concentrations of 1.9% or 3.7% by weight maintained their particle size (**Fig. sup2C**). Overall, adding the mucoadhesive agents changed the zeta potential value of the AS01_B_/OVA vaccine (**Fig. sup2B, D**), which was not surprising as all mucoadhesive agents tested were expected to have a certain extent of ionization at pH 7.2 to 7.4. However, subjecting the AS01_B_/OVA vaccine formulations to thin-film freeze-drying did not significantly affect their zeta potential values (**Fig. sup2B, D**). Shown in **Fig. 2A, B, C** are the particle sizes, particle size distribution curves, and zeta potential values of the AS01_B_/OVA vaccine formulations with CMC at 1.9%, 3.7%, 7.2%, or 16.3%, w/w, before and after they subjected to thin-film freeze-drying and reconstitution. Clearly, the particle size distribution profiles in the vaccine formulations containing 1.9% or 3.7% (w/w) of CMC were preserved after they were subjected to thin-film freeze-drying. The zeta potential values of the AS01_B_/OVA vaccines with CMC did not significantly change after they were subjected to thin-film freeze-drying (**Fig. 2C**), although it is unclear why the zeta potential value of the AS01_B_/OVA vaccine without CMC became more negative (**Fig. 2C**). Therefore, the AS01_B_/OVA vaccine formulations with CMC at 1.9% or 3.7% (w/w) were chosen for further study.

**Fig. 2.**
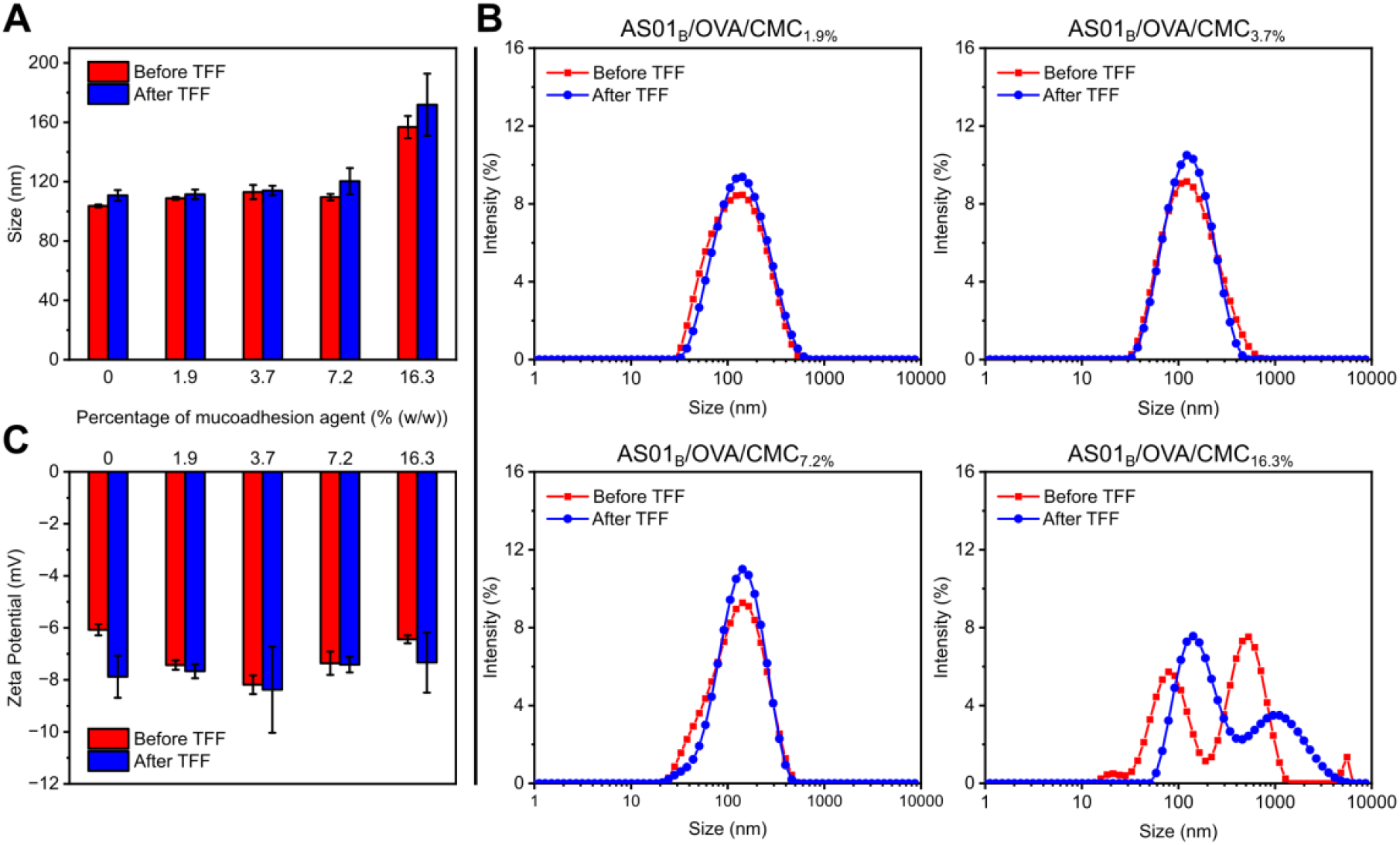
AS01_B_/OVA with different concentrations of CMC as a mucoadhesive agent. Shown are (A) particle size, (B) representative particle size distribution curve, and (C) zeta potential. Data in A and C are mean ± S.D. (n = 3).

### 3.3. The integrity of the OVA before and after the AS01_B_/OVA vaccine formulations with CMC were subjected to thin-film freeze-drying

To evaluate the impact of the thin-film freeze-drying process on the integrity of OVA antigen, SDS-PAGE was used to analyze the AS01_B_/OVA vaccine formulations containing 0, 1.9%, or 3.7% of CMC. The original OVA in solution (**Fig. 3**, lane I) showed two bands around 43 kDa, which might be attributed to the different glycosylated forms of OVA. A close comparison of **Fig. 3** lanes II vs. III, lanes IV vs. V, and lanes VI vs. VII did not show any apparent difference, indicating that the integrity of OVA was maintained, regardless of the content of the CMC in the vaccine formulations. SDS-PAGE analysis with the individual component of the AS01_B_/OVA vaccine formulations confirmed that the broad bands below 11 kDa shown in the AS01_B_/OVA samples (**Fig. 3**) were from the liposomes (**Fig. sup3**).

**Fig. 3.**
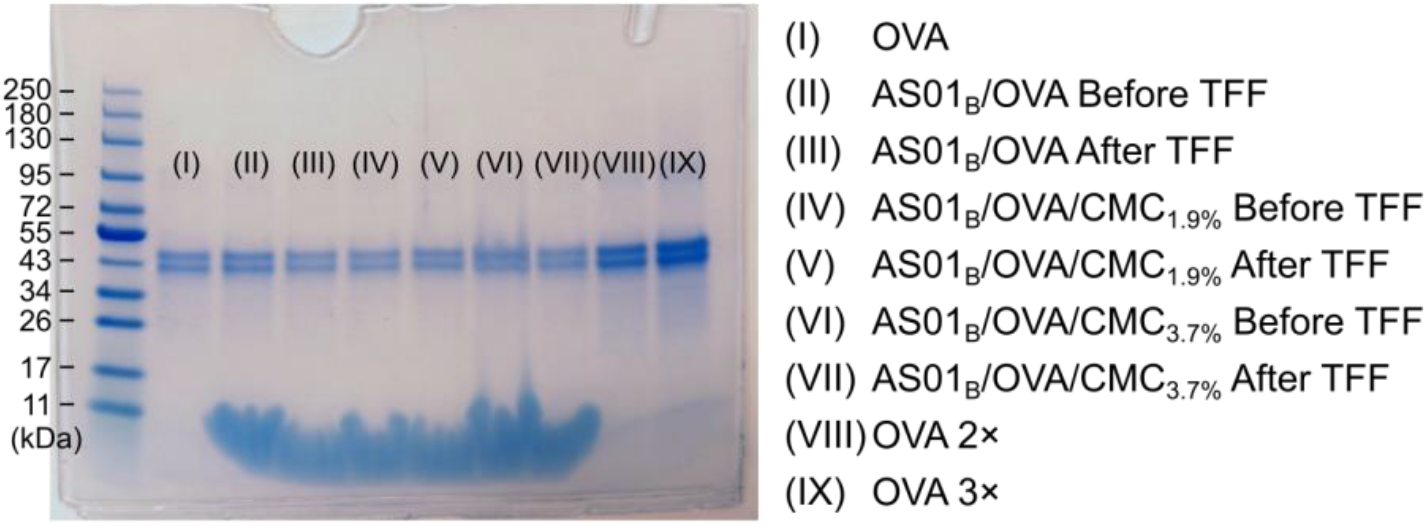
SDS-PAGE image of AS01_B_/OVA with 0, 1.9%, or 3.7% of CMC before and after they were subjected to thin-film freeze-drying and reconstitution.

### 3.4. In vitro mucoadhesion test

Petri dishes coated with 1.5% agar or 1.5% agar plus 2% porcine mucin were used to simulate the surface of human mucosal tissue. In the in vitro mucoadhesion test, films of TFF AS01_B_/OVA powders with 0, 1.9%, or 3.7% of CMC were placed on the agar or agar plus porcine mucin layer. The dishes were turned vertically and incubated at 32°C (**Fig. 4A**). The thin films swelled due to the moisture in the gel or the atmosphere and flowed down slowly. The maximum traveling distances of the formulations at different time points are shown in **Fig. 4B**.

**Fig. 4.**
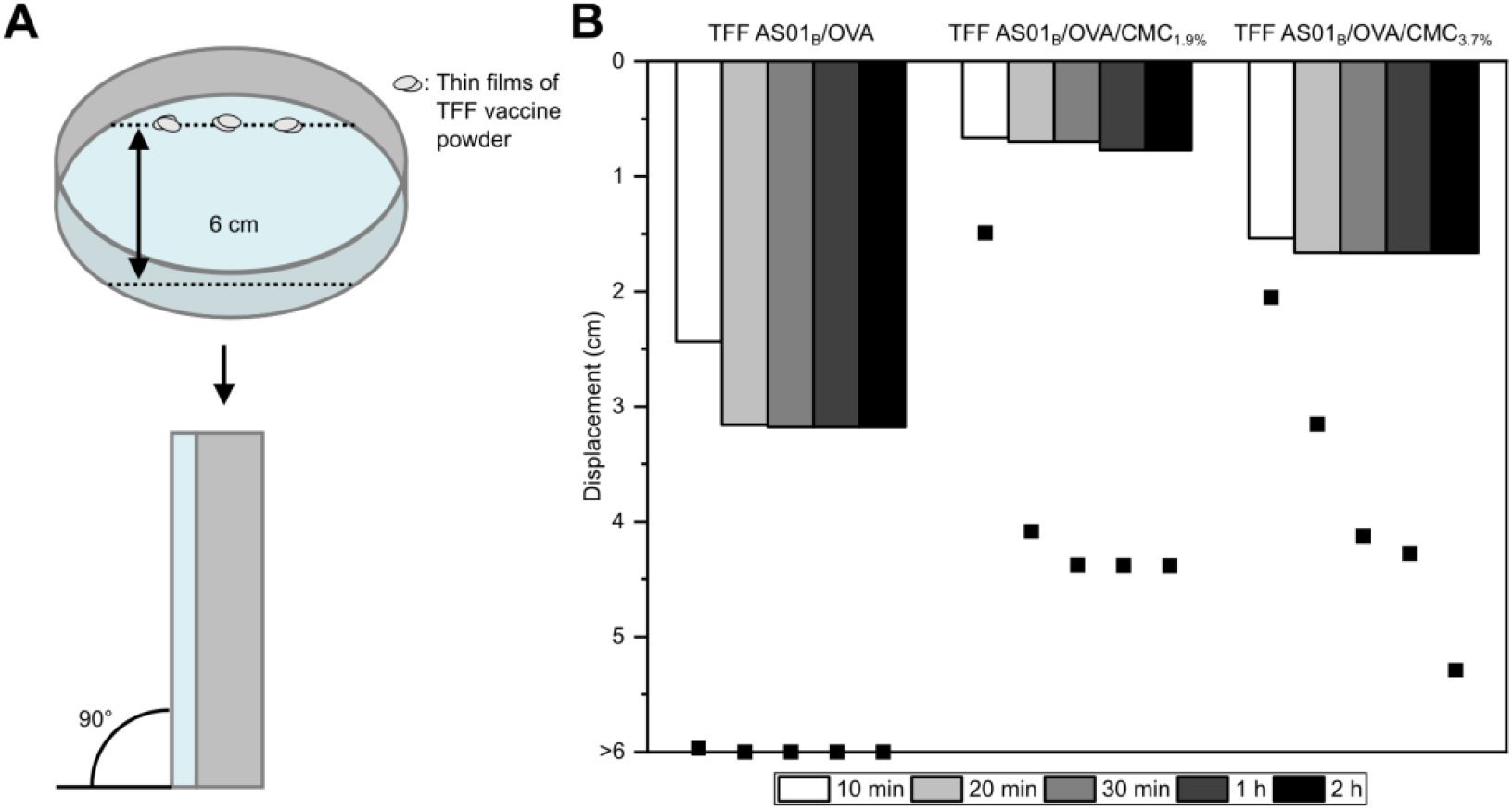
In vitro mucoadhesion test. (A) Experimental design. (B) Displacement of TFF AS01_B_/OVA vaccine powders with or without 1.9% or 3.7% CMC on 1.5% agar gel (dots) or 1.5% agar plus 2% porcine mucin gel (bars).

With the gel containing 1.5% agar only, the displacement of the TFF AS01_B_/OVA powder almost reached 6 cm after 10 min of incubations, while the displacements of TFF AS01_B_/OVA vaccine powders with 1.9% or 3.7% CMC were only 1.5 cm and 2 cm, respectively, during the same time period. The displacement of the TFF AS01_B_/OVA vaccine powders with 1.9% and 3.7% CMC both reached ∼4.5 cm after 1 h of incubation, indicating that the CMC in the AS01_B_/OVA vaccine rendered the vaccine powders adhesive to agar gel.

With the gel containing 1.5% agar and 2% porcine mucin, the displacements of all three TFF vaccine powders decreased significantly. However, the displacement of the TFF AS01_B_/OVA powder without CMC was still larger than that of TFF AS01_B_/OVA powders with 1.9% or 3.7% CMC (**Fig. 4B**). The displacement of the TFF AS01_B_/OVA powder with 1.9% CMC was slightly smaller than that with 3.7% CMC. Therefore, the TFF AS01_B_/OVA vaccine with 1.9% CMC (AS01_B_/OVA/CMC_1.9%_) was selected for further study.

### 3.5. Characterization of the TFF AS01_B_/OVA vaccine powders with 0 or 1.9% CMC

The residue water content in the TFF vaccine powders is important to their stability and aerosol properties. The moisture contents in the TFF AS01_B_/OVA powder and the TFF AS01_B_/OVA/CMC_1.9%_ powder were 3.29 ± 0.21% and 1.78 ± 0.14%, respectively. The CMC in the TFF AS01_B_/OVA/CMC_1.9%_ sample likely made the sucrose matrix less hygroscopic.

Surface profilometer was used to characterize the surface of the TFF vaccine powders. **Fig. 5A and 5B** show the images of the thin films of TFF AS01_B_/OVA powders with 0 or 1.9% of CMC. It appears that the TFF AS01_B_/OVA film was more porous compared to the TFF AS01_B_/OVA/CMC_1.9%_ film. In addition, the TFF AS01_B_/OVA/CMC_1.9%_ film had a radial pattern, which was likely because the nucleation process of the sucrose was altered by the CMC during the TFF process. The mean thickness of the TFF films was determined using six randomly selected films of the TFF AS01_B_/OVA vaccine powders with 0 or 1.9% CMC. The mean thickness of the TFF AS01_B_/OVA/CMC_1.9%_ films was 228.7 ± 51.1 μm, about 37% larger than that of the TFF AS01_B_/OVA films (i.e., 167.3 ± 44.4 μm). Considering that the distance between the needle tip and the cryogenically cooled surface was fixed, it is likely that the CMC in the AS01_B_/OVA vaccine made the vaccine droplets difficult to completely spread upon impact on the cooled surface during the TFF process. It is also possible that for the AS01_B_/OVA vaccine, without CMC, the sucrose was less prone to collapse during the drying process.

**Fig. 5.**
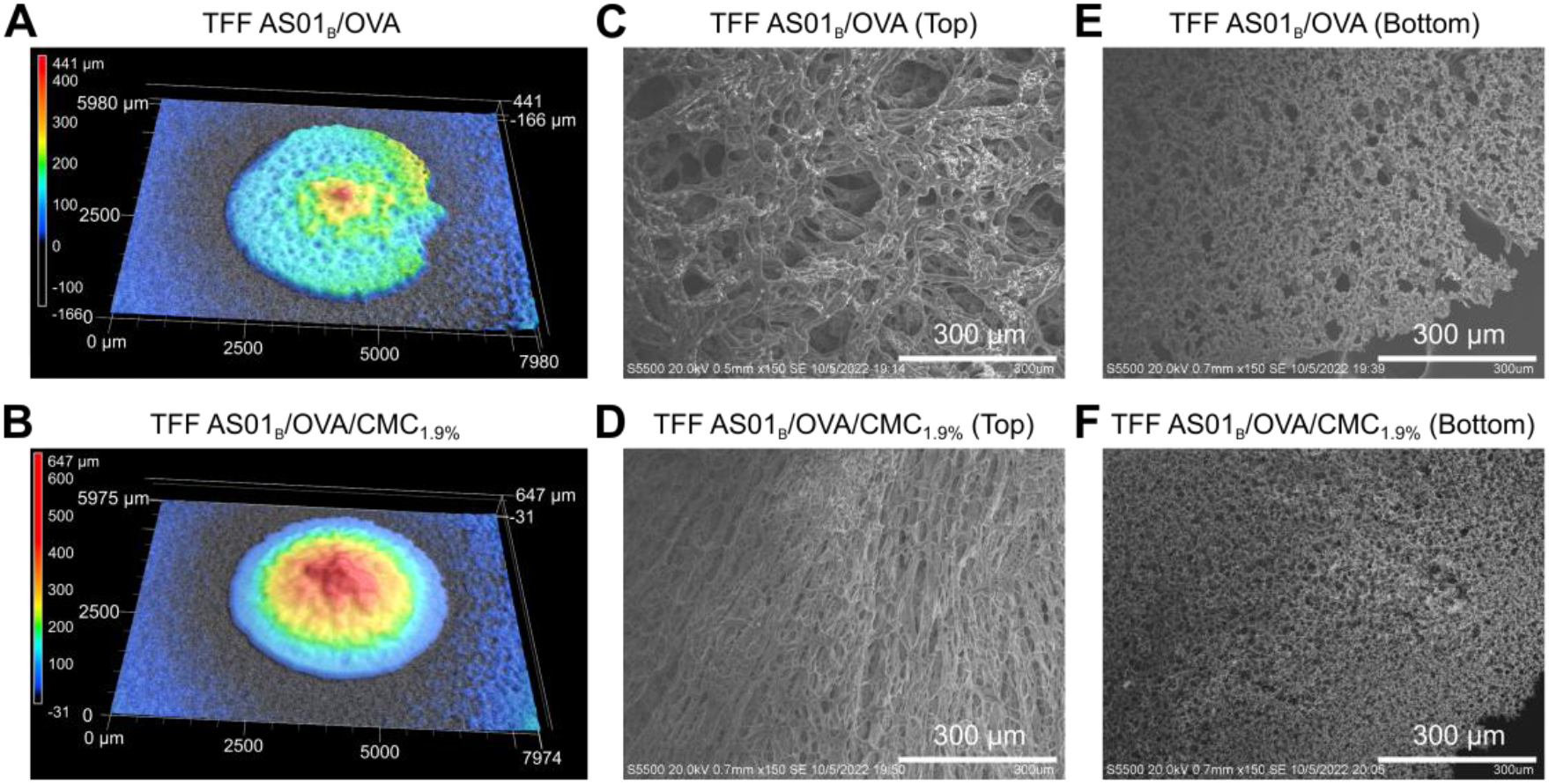
Characterization of TFF AS01_B_/OVA and AS01_B_/OVA/CMC_1.9%_ films. Shown in A-B are representative surface profiles of the TFF AS01_B_/OVA (A) and TFF AS01_B_/OVA/CMC_1.9%_ films (B). (C-D) Representative SEM images (top view) of the TFF AS01_B_/OVA (C) and AS01_B_/OVA/CMC_1.9%_ films (D). (E-F) Representative SEM images (bottom view) of the TFF AS01_B_/OVA (E) and AS01_B_/OVA/CMC_1.9%_ films (F).

The microscopic structures of the TFF vaccine powders were examined using SEM. The SEM images (**Fig. 5C-F**) (both top view and bottom view) revealed that the TFF AS01_B_/OVA and AS01_B_/OVA/CMC_1.9%_ films were porous. However, the TFF AS01_B_/OVA/CMC_1.9%_ sample looked more fibrous than the TFF AS01_B_/OVA sample (**Fig. 5C-D**), likely due to the presence of the CMC in the formulation. Moreover, minor phase separation was observed in both samples. The small particles deposited on the sucrose matrix were likely crystals of the buffer salts.

One important factor affecting the integrity of the vaccine formulation during the freezing and subsequent sublimation process was the crystallization of the excipients. XRD was used to characterize the crystalline properties of TFF powders (**Fig. 6A**). The peaks at 27.5°, 31.8°, 45.6°, 56.7°, 66.5°, and 75.6 on the patterns of TFF AS01_B_/OVA and TFF AS01_B_/OVA/CMC_1.9%_ powders were identified as the (111), (200), (220), (222), (400), and (420) planes of NaCl according to its simulated pattern. The crystallite sizes of the NaCl in TFF AS01_B_/OVA and AS01_B_/OVA/CMC_1.9%_ powders were 23.0 nm and 22.5 nm, respectively, as calculated using the Scherrer equation. Peaks unique to sucrose were not present, indicating that sucrose remained amorphous in the TFF vaccine powders, which is in agreement with previous findings with other sucrose-containing TFF powders (AboulFotouh et al., 2022b; Zhang et al., 2021), and is desirable for maintaining the integrity of the vaccine during TFF process.

**Fig. 6.**
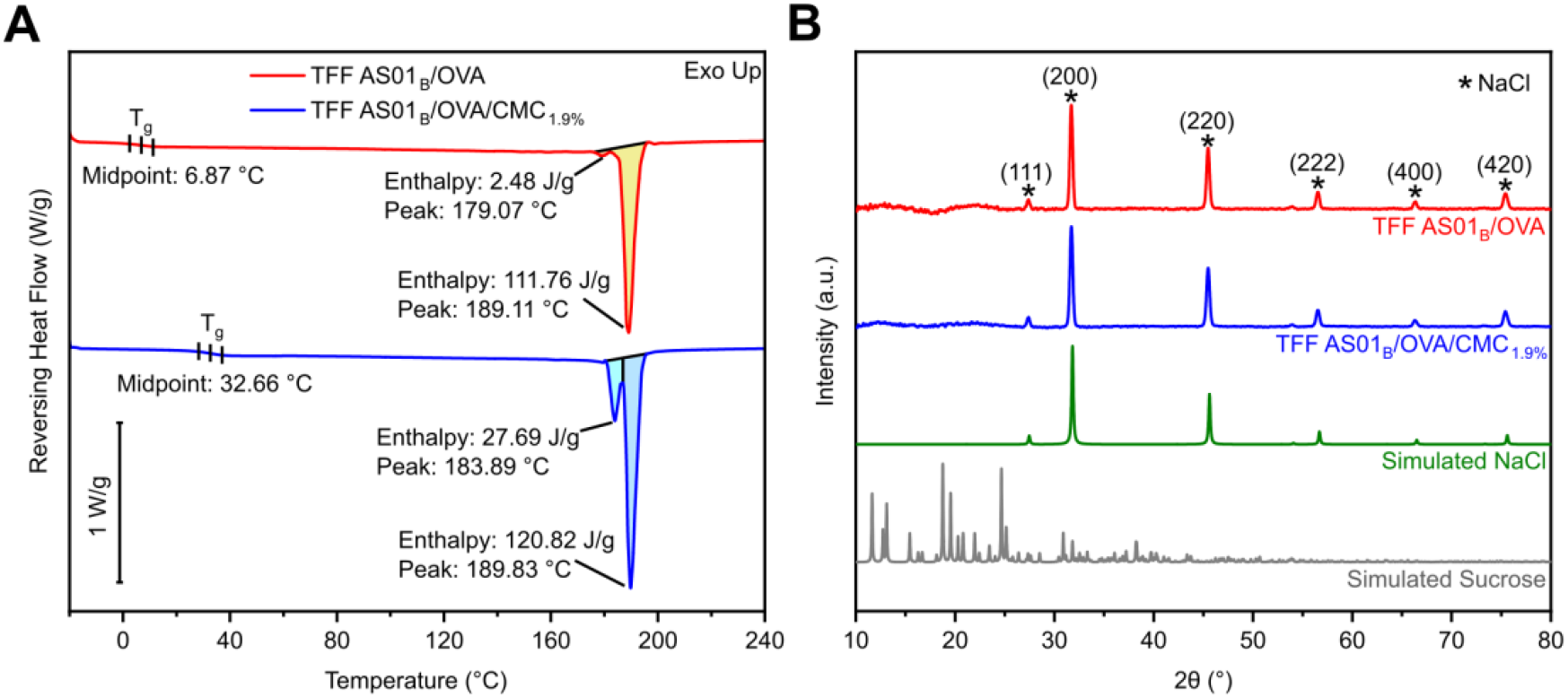
(A) The XRD diffractograms of the TFF vaccine powders and the simulated patterns of NaCl (COD: 1000041) and sucrose (COD: 3500015). (B) The DSC profiles of the TFF vaccine powders.

**Fig. 6B** shows the DSC profiles of the TFF AS01_B_/OVA and AS01_B_/OVA/CMC_1.9%_ samples. Their glass transition temperatures (T_g_) were 6.87°C and 32.66°C, respectively. Because it was reported that freeze-dried samples would crystalize and collapse when being stored at a temperature above their T_g_ (te Booy et al., 1992), it was concluded that the TFF AS01_B_/OVA/CMC_1.9%_ powder could provide better stability while stored at a temperature equal to or below room temperatures. Of course, different sugars and polymers may be included in the powders to further increase their T_g_, and thus thermostability, if needed. The DSC profiles of TFF AS01_B_/OVA and AS01_B_/OVA/CMC_1.9%_ powders both showed a major endotherm peak around 189°C, which is likely due to the melting of sucrose (Beckett et al., 2006). The mechanism underlying the smaller peak located left of the main melting endotherm of sucrose in both TFF AS01_B_/OVA and AS01_B_/OVA/CMC_1.9%_ powders is unknown. However, considering that there were several components in the TFF powders and sucrose is prone to decomposition during the melting process (Lu et al., 2017), the melting endotherm of sucrose could be distorted (Bhandari and Hartel, 2002).

### 3.6. Spraytec analysis of the thin-film freeze-dried AS01_B_/OVA vaccine powders sprayed with a UDSP nasal device

The AS01_B_/OVA vaccine powder was filled into the UDSP nasal device manually (∼19 mg per device) and sprayed to characterize the particle size distribution using laser diffraction. The existing FDA’s draft guidelines for size distribution characterization by laser diffraction only cover sprayed liquid formulations. The spray is divided into the initial phase, the fully developed phase, and the dissipation phase. According to the transmission profile shown in **Fig. 7A**, the TFF AS01_B_/OVA vaccine powder sprayed with the UDSP nasal device did not have an obvious fully developed (or stable) phase. Instead, the transmission reached a minimum around 3 milliseconds (ms) after the actuation and then rapidly recovered to the maximum within 14 ms; this result matched the time-course images of the sprayed powder captured by a high-speed camera (**Fig. 7B**). Therefore, the particle size at 4 ms centered around the minimum transmission was integrated, which was more representative compared to that in the recommended fully developed phase.

**Fig. 7.**
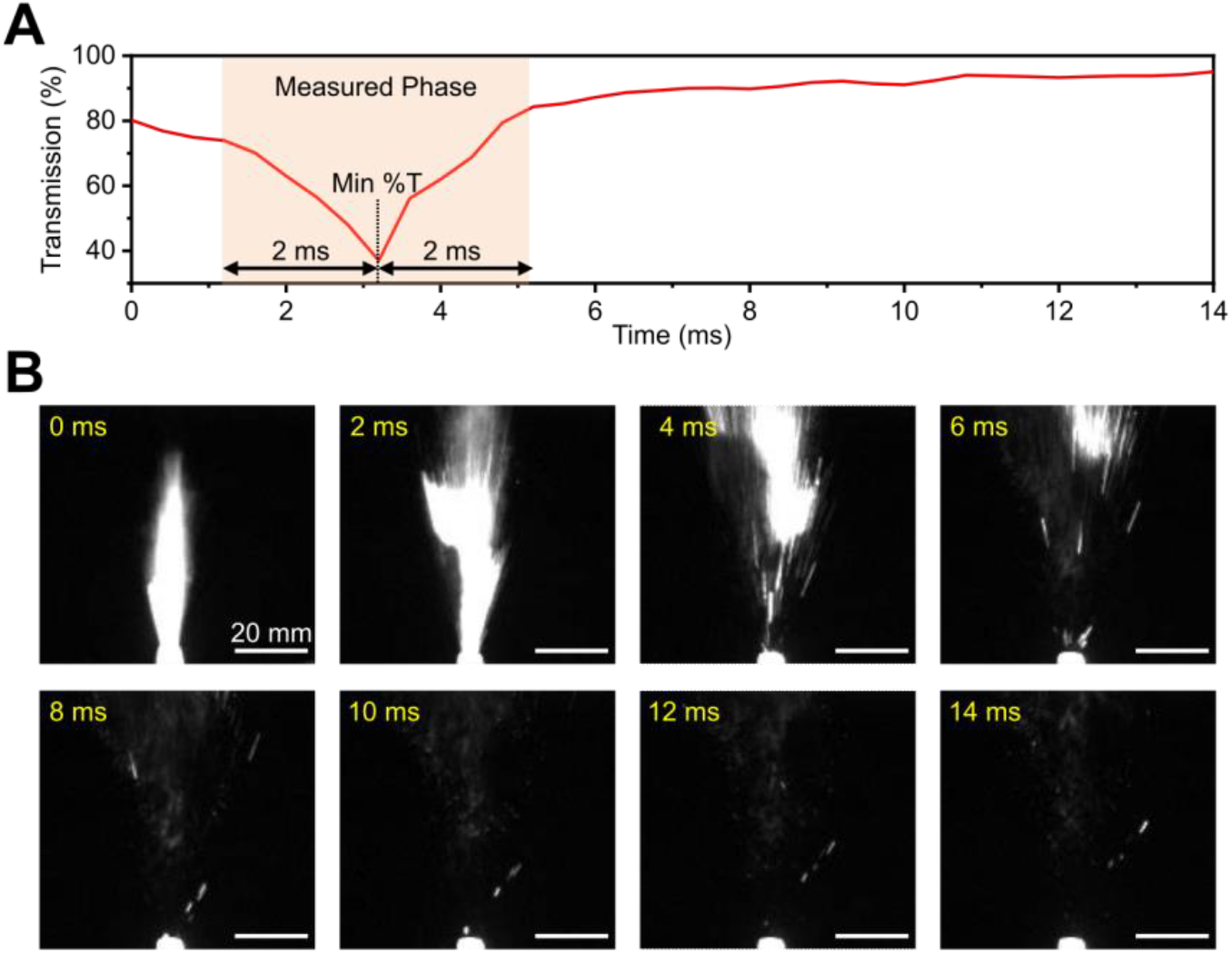
(A) A representative transmission profile of the TFF AS01_B_/OVA vaccine powder and (B) the time course images of the TFF AS01_B_/OVA powder sprayed with the UDSP nasal device.

**Figs. 8A** and **B** show the particle size distribution of the TFF AS01_B_/OVA and the TFF AS01_B_/OVA/CMC_1.9%_ powders, which had a median particle size by volume (Dv(50)) value of 311.9 ± 1.3 μm and 224.2 ± 12.0 μm, respectively. The %V < 10 μm values for the TFF AS01_B_/OVA powder and the AS01_B_/OVA/CMC_1.9%_ powder were 0.66 ± 0.09 % and 0.51 ± 0.09 %, respectively (**Fig. 8A, B**).

**Fig. 8.**
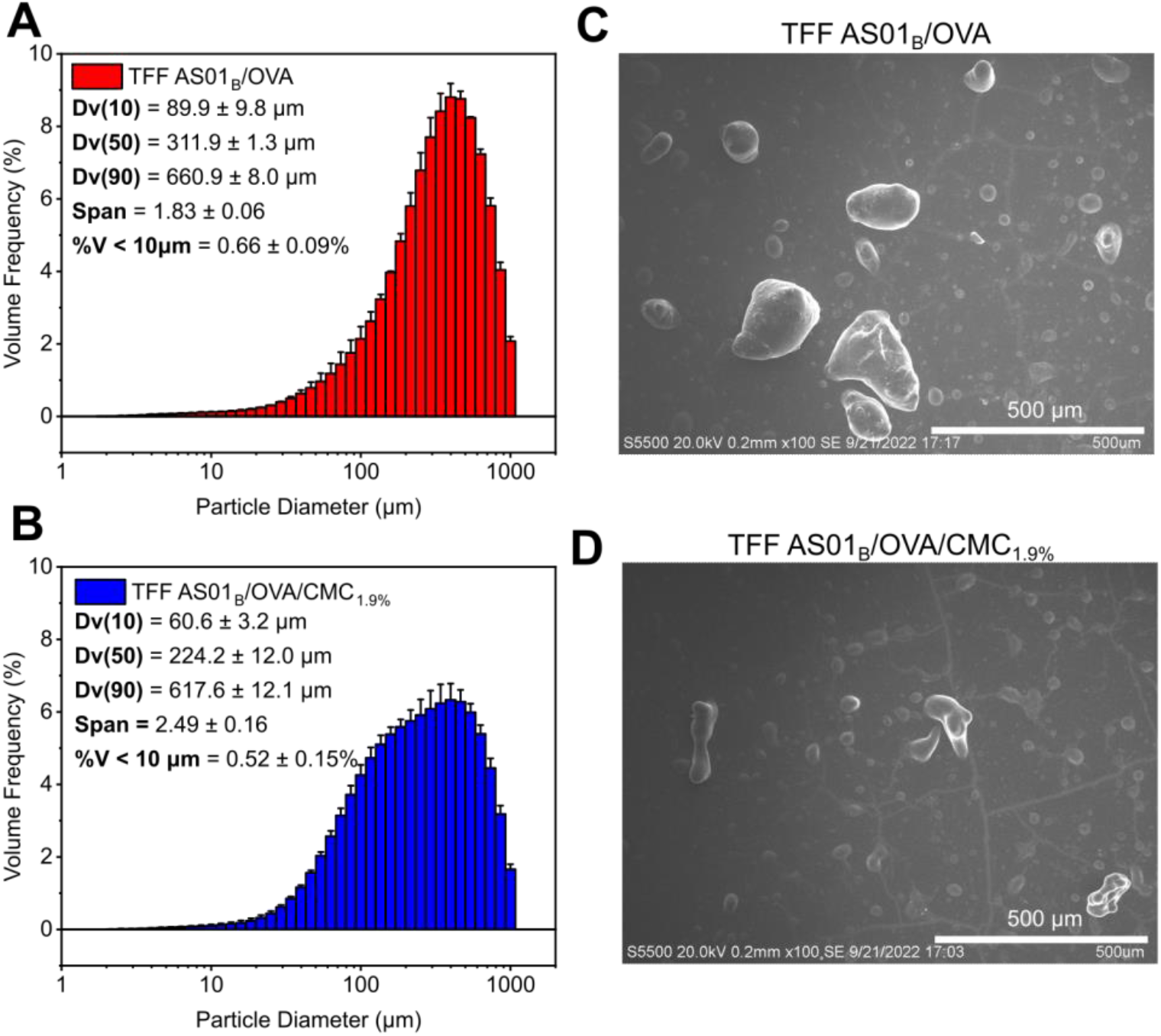
(A-B) The particle size distribution curves of (A) TFF AS01_B_/OVA and (B) TFF AS01_B_/OVA/CMC_1.9%_ powders sprayed using a UDSP nasal device and determined using a Spraytec spray particle size analyzer. (**C-D**) Representative SEM images of TFF AS01_B_/OVA powder and TFF AS01_B_/OVA/CMC_1.9%_ powder after they were sprayed using a UDSP nasal device.

SEM was used to examine the sprayed powders of the TFF AS01_B_/OVA and the AS01_B_/OVA/CMC_1.9%_ vaccines (**Fig. 8C, D**). The particles of sprayed powders were not uniform, showing a wide size distribution, and the particles of the sprayed TFF AS01_B_/OVA/CMC_1.9%_ powder were generally smaller than that of sprayed TFF AS01_B_/OVA powder, which agreed with the Spraytec results (**Fig. 8A, B**). Although there were small particles in the background of sprayed TFF AS01_B_/OVA powder and AS01_B_/OVA/CMC_1.9%_ powder (**Fig. 8C, D**), the contribution of those particles to the volume frequency was insignificant compared to large particles, which agrees with the unimodal distribution of the particle size based on the Spraytec result (**Fig. 8A, B**). Also, since most of the small particles were larger than 10 μ;m, the risk of the particles entering the lung after intranasal administration in humans is minimized, indicating that it is suitable to use the UDSP nasal device for intranasal administration of the TFF vaccine powders.

### 3.7. The plume geometry and spray pattern of the thin-film freeze-dried AS01_B_/OVA vaccine powders sprayed with the UDSP nasal device

The plume geometry and the spray pattern of the TFF AS01B/OVA vaccine powder were studied after the powder was sprayed using a UDSP nasal device and shown in **Fig 9**. The plume angles of the TFF AS01_B_/OVA powder and the TFF AS01_B_/OVA/CMC_1.9%_ powder were 24.90 ± 4.05° and 24.52 ± 4.81°, respectively. Statistical analysis of the spray pattern shows that the area of the pattern was 367.70 ± 48.96 mm^2^ for the TFF AS01_B_/OVA powder and 361.70 ± 55.83 mm^2^ for the TFF AS01_B_/OVA/CMC_1.9%_ powder. Both powders had good ovality values, i.e., 1.28 ± 0.11 for the TFF AS01_B_/OVA powder and 1.22 ± 0.08 for the TFF AS01_B_/OVA/CMC_1.9%_ powder. Overall, it was concluded that including the CMC at 1.9% w/w in the AS01_B_/OVA vaccine did not significantly affect the spray pattern and plume geometry of the powder after spraying using the UDSP nasal device.

**Fig. 9.**
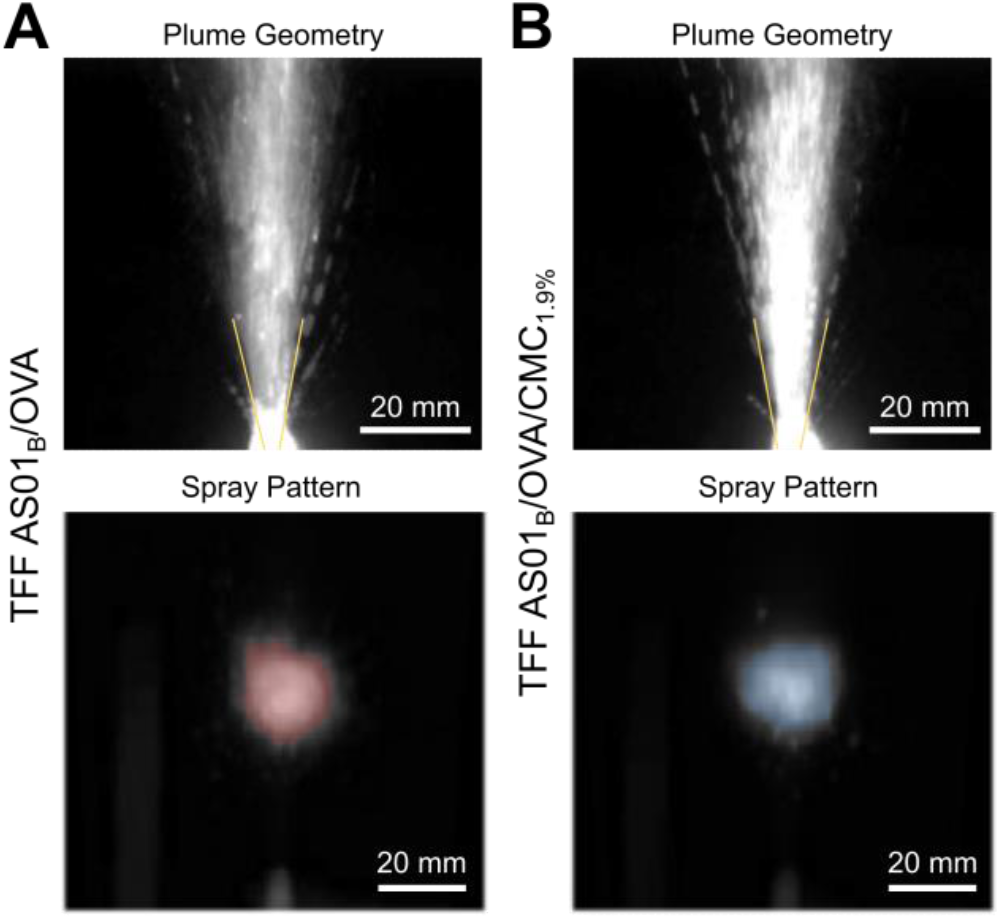
The plume geometries and spray patterns of the TFF AS01_B_/OVA and the TFF AS01_B_/OVA/CMC_1.9%_ powders after they were sprayed using a UDSP nasal device.

### 3.8. The integrity of the OVA and the AS01_B_ after the AS01_B_/OVA vaccine powders were sprayed using the UDSP nasal device

The integrity of the OVA after the TFF AS01_B_/OVA powder was aerosolized using the UDSP nasal device was evaluated using SDS-PAGE. As shown in **Fig. 10A**, a comparison of the OVA bands in lanes II vs. III and lanes IV vs. V did not reveal fragmentation or aggregation of the OVA protein in the TFF AS01_B_/OVA powder and the TFF AS01_B_/OVA/CMC_1.9%_ powder. Clearly, the OVA protein in the AS01_B_/OVA vaccine powders was not sensitive to the shear stress associated with spraying them with the UDSP nasal device.

**Fig. 10.**
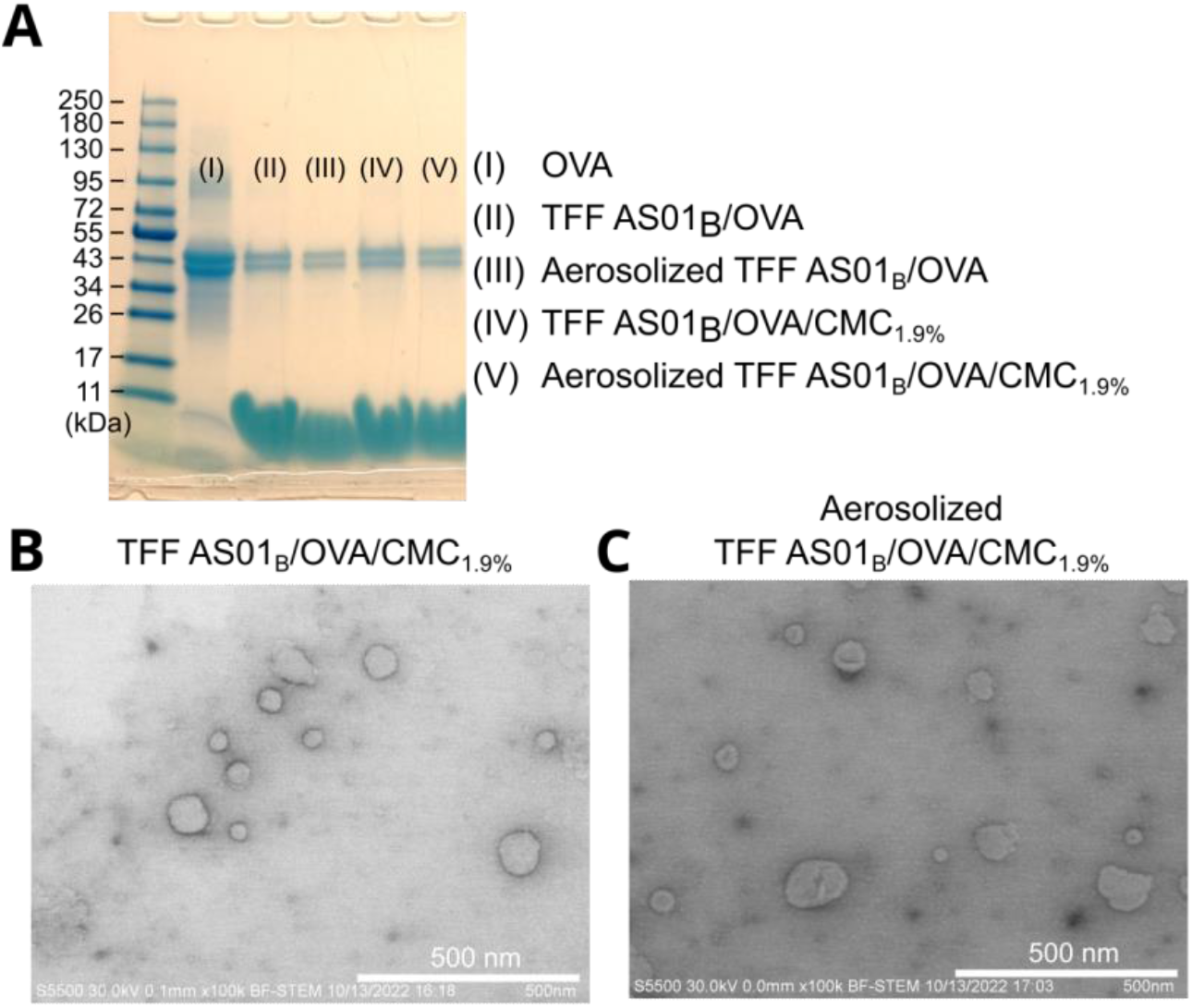
The integrity of the AS01_B_/OVA vaccine powder before and after being sprayed with the UDSP nasal device. (A) The SDS-PAGE image of the TFF AS01_B_/OVA and the AS01_B_/OVA/CMC_1.9%_ before and after they were sprayed. (B) Representative STEM images of TFF AS01_B_/OVA/CMC_1.9%_ powder before and after it was sprayed using a UDSP device.

STEM was also used to characterize the TFF AS01_B_/OVA/CMC_1.9%_ powder before and after it was sprayed with the UDSP nasal device (**Fig. 10B, C**). The liposomes in the TFF AS01_B_/OVA/CMC_1.9%_ powder before and after being sprayed with the UDSP nasal device were about 100 nm. Also, no obvious liposome aggregation was observed after the TFF AS01_B_/OVA/CMC_1.9%_ powder was sprayed, indicating that the liposomes in the TFF AS01_B_/OVA/CMC_1.9%_ vaccine powders were also not sensitive to the shear stress associated with spraying them with the UDSP nasal device.

### 3.9. *Deposition patterns of the TFF* AS01_B_/OVA/CMC_1.9%_ *vaccine powder in nasal casts*

The TFF AS01_B_/OVA/CMC_1.9%_ powder was manually loaded into the UDSP nasal device at 22.99 ± 1.58 mg (n = 27). After actuation, the delivered amount of the TFF AS01_B_/OVA/CMC_1.9%_ powder was 21.69 ± 1.60 mg. In other words, the delivery efficiency was 94.46 ± 4.87%.

The deposition pattern of the TFF AS01_B_/OVA/CMC_1.9%_ in the nasal cast based on the CT scan of a 48-year-old male is shown in **Fig. 11A-B**. In all the conditions, the majority of the TFF powder was deposited in the middle and lower turbinate and the nasopharynx regions. When an air flow of 10 LPM was applied to the nasal cast, the deposition percentage in the nasopharynx region and the filter increased, which is expected. The percentage deposited in the upper turbinate region where the olfactory bulb resides was less than 10% in all the conditions, indicating that the exposure of the vaccine powder to the olfactory bulb can be minimized by spraying the TFF vaccine powder with the UDSP nasal device. As shown in **Fig. 11C**, the RDP of Run #1 had the highest value of 84.4%, and the Run #2 showed the lowest value of 71.3%. To evaluate the effect of each parameter on the RDP, the S/N ratios were calculated. Since a higher S/N ratio will lead to a higher RDP, it was concluded that the optimal condition for the RDP in the nasal cast based on a 48-year-old male is: 0° for the coronal angle, 45° for the sagittal angle, and 0 LPM for the flow rate, which is the same as Run #1 (**Fig. 11D**). Finally, since the difference of the S/N ratio between two levels indicates the impact of the parameter on the RDP, it was concluded that the order of the impact on RDP was sagittal angle > coronal angle > flow rate.

**Fig. 11.**
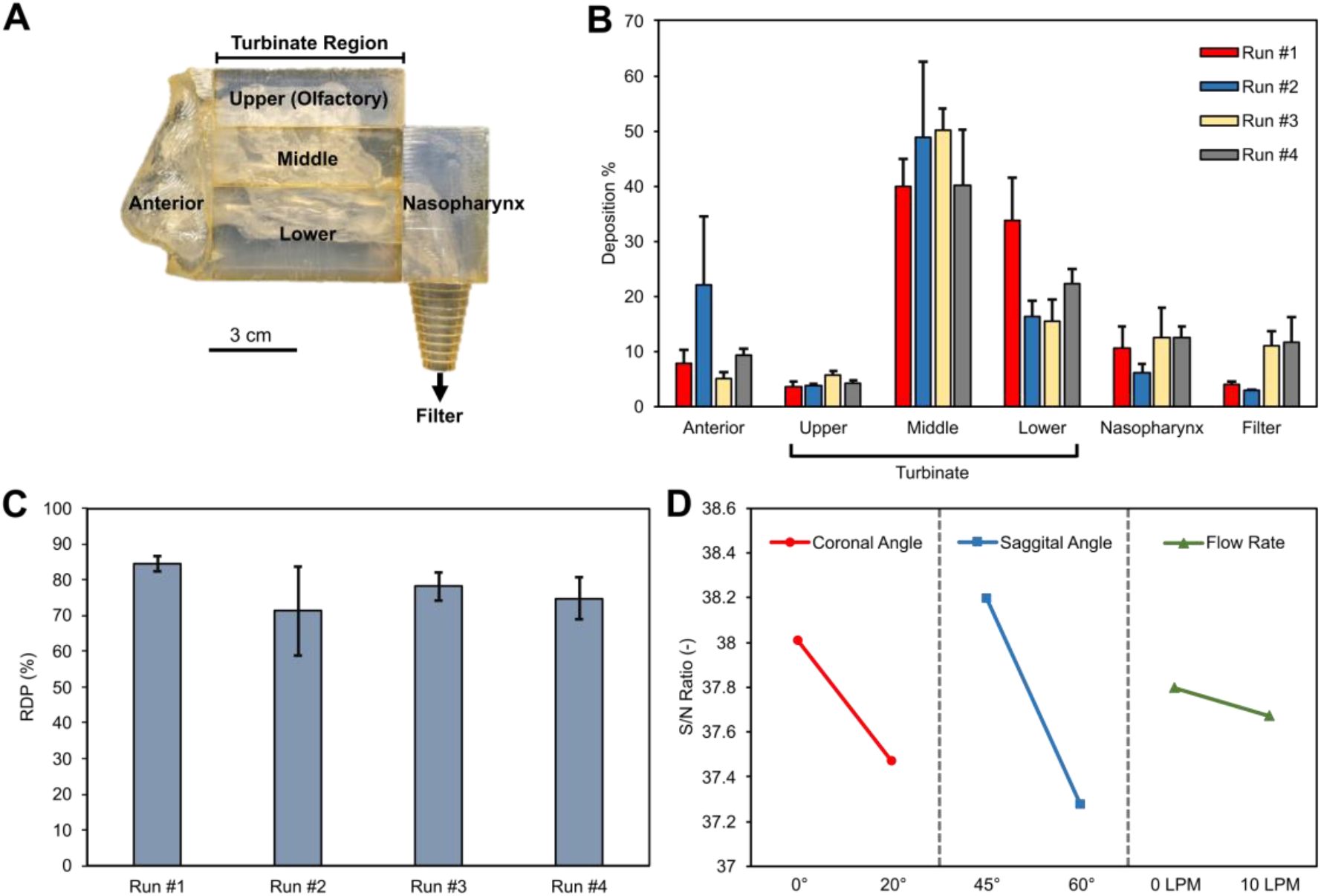
Deposition patterns of TFF AS01_B_/OVA/CMC_1.9%_ vaccine powder in a nasal cast based on a 48-year-old male. (A) A digital image of the nasal cast with different regions labeled. (B) The deposition patterns of the TFF AS01_B_/OVA/CMC_1.9%_ powder in different regions. The parameters were designed with the Taguchi array method. (C) The regional deposition efficiency (RDP) of the 4 conditions in the middle and lower turbinate and the nasopharynx regions. (D) The S/N ratio of each parameter at different levels.

The deposition pattern of the TFF AS01_B_/OVA/CMC_1.9%_ powder in a nasal cast based on the CT scan of a 7-year-old female is shown in **Fig. 12A-B**. The result showed that most of the TFF AS01_B_/OVA/CMC_1.9%_ powder was also deposited in the middle and lower turbinate regions and the nasopharynx region. Compared to the results generated using the adult nasal cast, the differences between different conditions were more significant in the child nasal cast. For example, Run #1 resulted in the highest deposition percentage in the nasopharynx region (**Fig. 12B**), which was desirable for intranasal immunization. This was likely due to the smaller size/depth of the nasal cavity in children, making the effect of these conditions more pronounced. The RDP among 4 conditions ranged from 75.8% to 88.8% (**Fig. 12C**), which were generally higher than that of the adult nasal cast. The calculated S/N ratios for all parameters indicated that the optimal condition for the deposition pattern was also: 0° for the coronal angle, 45° for the sagittal angle, and 0 LPM for the flow rate, which is the same as Run #1 (**Fig. 12D**). Different from the adult nasal cast, however, the order of the impact on RDP was: coronal angle > sagittal angle > flow rate.

**Fig. 12.**
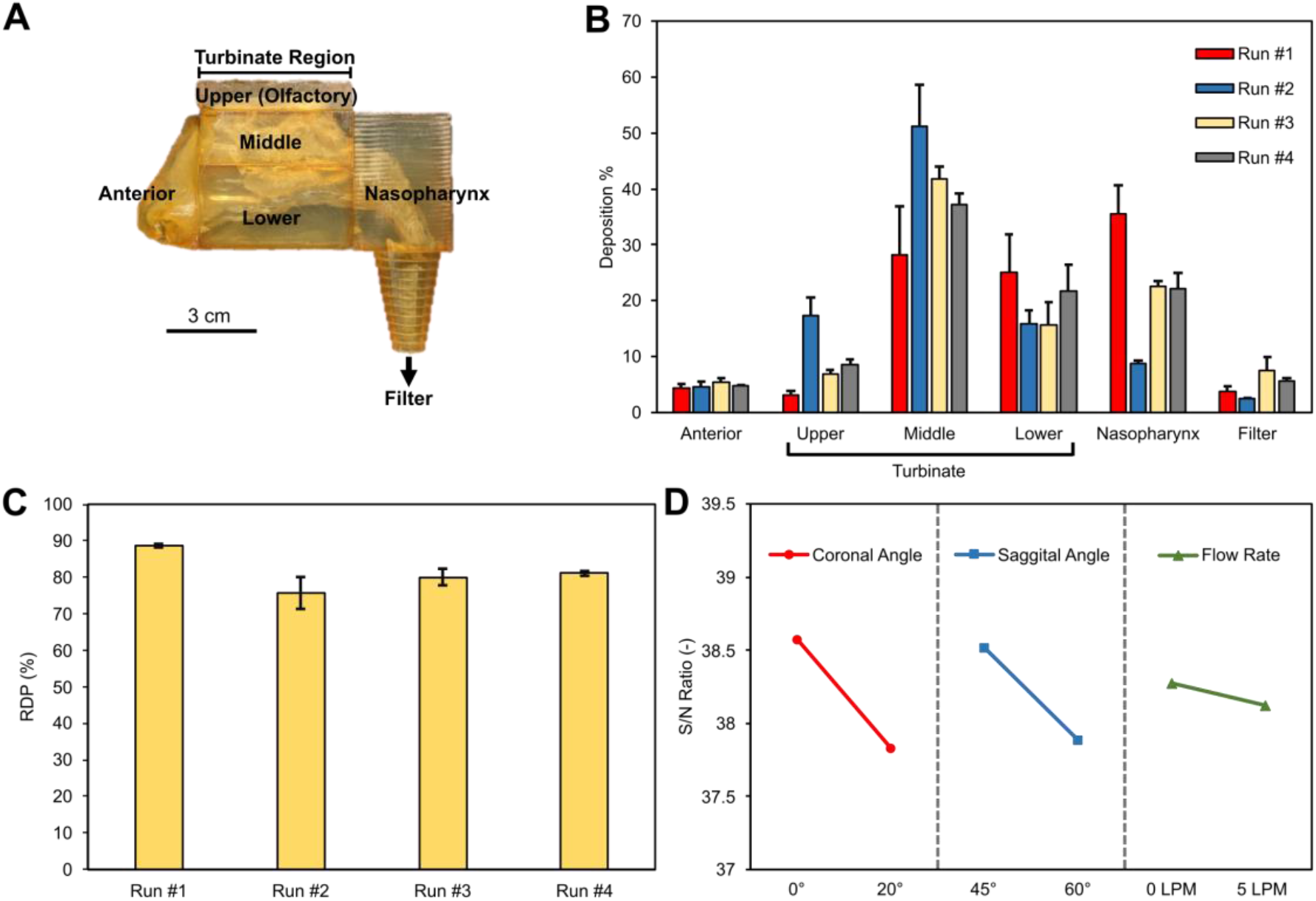
Deposition pattern of TFF AS01_B_/OVA/CMC_1.9%_ vaccine powder in a nasal cast based on a 7-year-old female. (A) A digital image of the nasal cast with different regions labeled. (B) The deposition patterns of the TFF AS01_B_/OVA/CMC_1.9%_ powder in different regions. The parameters were designed with the Taguchi array method. (C) The regional deposition efficiency (RDP) of the 4 conditions in the middle and lower turbinate and the nasopharynx regions. (D) The S/N ratio of each parameter at different levels.

Taken together, data in **Fig. 11** and **Fig. 12** showed that in both adult and child nasal casts, the optimal condition to deliver the TFF AS01_B_/OVA/CMC_1.9%_ vaccine powder to the middle and lower turbinate and the nasopharynx region was 0° for the coronal angle, 45° for the sagittal angle, and 0 LPM for the flow rate. The insertion depth is expected to also significantly affect the deposition pattern of the TFF vaccine powder sprayed using the UDSP nasal device (Gao et al., 2020). In the present study, for the adult nasal cast, the insertion depth was 0.5 cm and 0.4 cm for the child nasal cast, so that all the parameters in the Taguchi L8 orthogonal array in **Table 1** can be accommodated. Due to the rigid structure of the 3D-printed nasal casts, the insertion depth could not be adjusted in a large interval. Further optimization of the insertion depth may be considered in the future.

**Fig. sup6A** showed the deposition percentages of the TFF AS01_B_/OVA/CMC_1.9%_vaccine powder directly in the nasopharynx region of a 48-year-old male nasal cast in 4 different runs. The N/S ratios in **Fig. sup6B** indicated that the optimal condition for the TFF vaccine powder to be delivered directly to the nasopharynx region of the nasal cast based on the 48-year-old male is: 0° for the coronal angle, 45° for the sagittal angle, and 10 LPM for the flow rate. The deposition pattern of the TFF AS01_B_/OVA/CMC_1.9%_ vaccine powder at the optimal condition identified above is shown in **Fig. sup7**. The percentage of the powder deposited directly in the nasopharynx region was 15.09 ± 4.05%.

**Fig. sup8A** showed the deposition percentages of the TFF AS01_B_/OVA/CMC_1.9%_vaccine powder directly in the nasopharynx region of the 7-year-old female nasal cast in 4 different runs. The N/S ratios in **Fig. sup8B** indicated that the optimal condition for the TFF vaccine powder to be delivered directly to the nasopharynx region of the nasal cast based on the 7-year-old female is 0° for the coronal angle, 45° for the sagittal angle, and 5 LPM for the flow rate. Even at 0° for the coronal angle, 45° for the sagittal angle, and 0 LPM flow rate, the percentage of the powder deposited directly in the nasopharynx region was 35.49 ± 5.14%. Therefore, if one only considers the TFF AS01_B_/OVA/CMC_1.9%_ vaccine powder that is directly delivered into the nasopharynx region, where the Waldeyer’s ring is located, then applying a flow rate, similar to applying the vaccine to a human subject while the human subject is inhaling, is expected to be beneficial, but there is also an increased chance for the powder to travel beyond the nasopharynx region into the trachea. Direct deposition of a vaccine to the nasopharynx region is important as the vaccine is expected to have a higher chance of reaching the lymphoid tissues (i.e., adenoids and tubal tonsils) in the Waldeyer’s ring. If the vaccine is deposited in the posterior nasal cavity, it may ultimately reach the naso-oropharynx region because of the mucous blanket posterior movement, but the lymphoid tissues in the naso-oropharynx need to efficiently interact with the vaccine or antigens bound to the mucus.

The results reported in this study showed that a protein antigen-based vaccine that is adjuvanted with the liposomal AS01_B_ adjuvant and contains CMC as a mucoadhesive agent could be converted into a dry powder using the TFF technology and then delivered to the desired regions in nasal casts 3D printed based on the CT-scan images of human noses using the UDSP nasal device. Besides the AS01_B_-adjuvanted vaccines, the TFF technology has been successfully applied to various other types of vaccines, including vaccines adjuvanted with aluminum salt (Alzhrani et al., 2021; Li et al., 2015; Thakkar et al., 2018), MF59 or AddaVax (AboulFotouh et al., 2022a), plasmid DNA vaccine (Xu et al., 2022), viral vector-based vaccines (Cui, unpublished data), and messenger RNA-lipid nanoparticle vaccines (Cui, unpublished data). In addition, both Gram-positive and Gram-negative bacteria in suspension can be converted to dry powders using the TFF technology (Wang et al., 2022). Therefore, it is expected that various other types of vaccines can be administered intranasally as TFF powders using the UDSP nasal device to target the vaccines to the lymphoid tissues in the nasopharynx region in humans. Of course, for different vaccines, additional optimization of the formulation and critical processing parameters will likely be needed.

For decades, there has been a strong interest in developing intranasal vaccines. However, as aforementioned, only a few nasal vaccines have received regulatory approval for human use around the world. Many nasal vaccine candidates that are safe and efficacious in preclinical studies often fail to move beyond phase 1 in clinical testing (Cai et al., 2022). For example, the ChAdOx1 nCoV-19 simian adenovirus-based COVID-19 vaccine that was effective in hamsters and non-human primates (NHP) when given intranasally failed to elicit consistent immune responses in humans (Madhavan et al., 2022). The authors hypothesized that the dosage form and the device used to administer the vaccine, both were not optimized for intranasal vaccination, may have contributed, at least in part, to the weak and inconsistent immune responses seen in humans (Madhavan et al., 2022), underscoring the significance of optimizing vaccine formulation and delivery device in nasal vaccine development.

Nasal vaccines that are currently approved for human use around the world are liquid suspension or freeze-dried powder that must be reconstituted before administration. They are sprayed into the nostrils using a nasal spray system such as the BD Accuspray™. Intranasal immunization using a vaccine dry powder is appealing, but a technology that can transform vaccines from liquid to a dry powder suitable for intranasal delivery, while preserving the physical and chemical properties and immunogenicity of the vaccines is needed. Similarly, a dry powder delivery system that can deliver the vaccine powder to the desired region(s) of the human nasal cavity is also required because there has not been an approved nasal dry powder vaccine for human use yet. Since many intranasal vaccines failed to move beyond phase 1 clinical trials (Cai et al., 2022; Xu et al., 2021b), the formulation of a vaccine intended for intranasal administration must be optimized, and only a device that can efficiently target the vaccine formulation to the desired region(s) in the nasal cavity should be chosen. As aforementioned, in children and adults, the Waldeyer’s ring in the naso-oropharynx region is the key lymphoid tissue in the nasal cavity. Therefore, it is ideal to deliver a vaccine powder directly to the nasopharynx region. If the vaccine is delivered in the posterior nasal cavity and binds to the mucus, then it may be carried to the naso-oropharynx region by the mucous blanket posterior movement by means of ciliary clearance, wherein the vaccine may be taken up by the lymphoid tissues in the Waldeyer’s ring or cleared through the throat to the stomach and then degraded or deactivated there (Sahin-Yilmaz and Naclerio, 2011). It is possible, or perhaps likely, that some cells in the epithelium of the posterior nasal cavity covered by the mucous layer may be able to take up the vaccine or the antigens, but the vaccine/antigen must penetrate through the mucous layer efficiently and quickly as the mucous blanket posterior movement can clear inhaled particles from the nasal cavity in 10-20 min (Sahin-Yilmaz and Naclerio, 2011).

A vaccine that is administered/deposited in the region that is anterior to the lower turbinate will be transported anterior out of the nostrils. Testing vaccine powders and delivery devices directly in human subjects is ideal, but less practical. Nasal casts 3D printed based on the CT scan images of the noses of humans of different ages, such as the nasal casts used in the present study and the idealized nasal replica (Chen et al., 2020), provide a suitable alternative for vaccine powder optimization and dry powder delivery device selection. Formulations and devices selected based on nasal casts can then be validated in humans to verify the deposition of the vaccine powder to desired regions or sites before initiating clinical trials to test the safety and efficacy of new vaccine candidates.

## 4. Conclusion

It is concluded that it is feasible to apply the TFF technology to convert vaccines such as the AS01_B_-adjuvanted OVA model vaccine with a proper mucoadhesive agent such as CMC into dry powders, and the resultant vaccine powders can be delivered to the desired regions of 3D printed human nasal casts using the UDSP nasal device. The type of the mucoadhesive agents and their concentration were critical so that the physical properties of the vaccine and the integrity of the antigen were preserved after the vaccine was subjected to the TFF technology. The mucoadhesive agent also affected the powder properties of the TFF vaccine powder. When sprayed using the UDSP nasal device, the TFF vaccine powders showed desirable particle size distribution, spray pattern and plum geometry. Importantly, spraying the vaccine powder with the UDSP nasal device did not negatively affect the antigen and the adjuvant. Finally, evaluation of the deposition patterns of TFF AS01_B_/OVA/CMC_1.9%_ powder in both adult and child nasal casts showed that the optimal parameters for both nasal casts were: 0° for the coronal angle, 45° for the sagittal angle, and 0 LPM for the flow rate, and more than 80% of the powders were deposited in the middle and lower turbinate and the nasopharynx regions. If only the nasopharynx region is considered, then applying a flow rate was beneficial.

## ACKNOWLEDGEMENTS

Z Cui and RO Williams III report financial support from TFF Pharmaceuticals, Inc. The UDSP device was generously provided by the AptarGroup, Inc. Y Yu was supported in part by the Y.L. Lin Hung Tai Education Foundation and the National Taiwan University. K AboulFotouh was supported in part by an Egyptian Government Fellowship. We thank Dr. Hugh Smyth in the College of Pharmacy at UT Austin for letting us use the laser sheet system in his laboratory to acquire the spray pattern and plume geometry data.

## CRediT Author Statement

Y Yu: Conceptualization, Methodology, Investigation, Visualization, Formal analysis, Writing Original Draft; K AboulFotouh: Methodology, investigation; G Williams, J Suman, C Cano: Conceptualization, Resources, Writing-Review & Editing; Z Warnken: Methodology, investigation, Writing-Review & Editing; RO Williams III: Resources, Writing-Review & Editing, Funding acquisition. Z Cui: Conceptualization, Resources, Validation, Writing-Review & Editing, Supervision, Funding acquisition.

## Supplementary Material

**S1. Preparation of the OVA-FITC**. OVA and (240 mg) and FITC (12 mg) were dissolved in 60 mL of 0.1 M sodium carbonate/bicarbonate solution (pH = 9). The mixture was stirred overnight at room temperature. After the reaction, the buffer and the unreacted FITC were removed by centrifugation with 30,000 MWCO centrifugal filter tubes from Millipore (Grand Island, NY, USA). The FITC-OVA was then washed 4 times with 20 mL of PBS. To purify the FITC-OVA, the mixture was further centrifuged with the 100,000 MWCO centrifugal filter tubes, and the filtrate (i.e., OVA-FITC in PBS) was collected.

**Fig. sup1.**
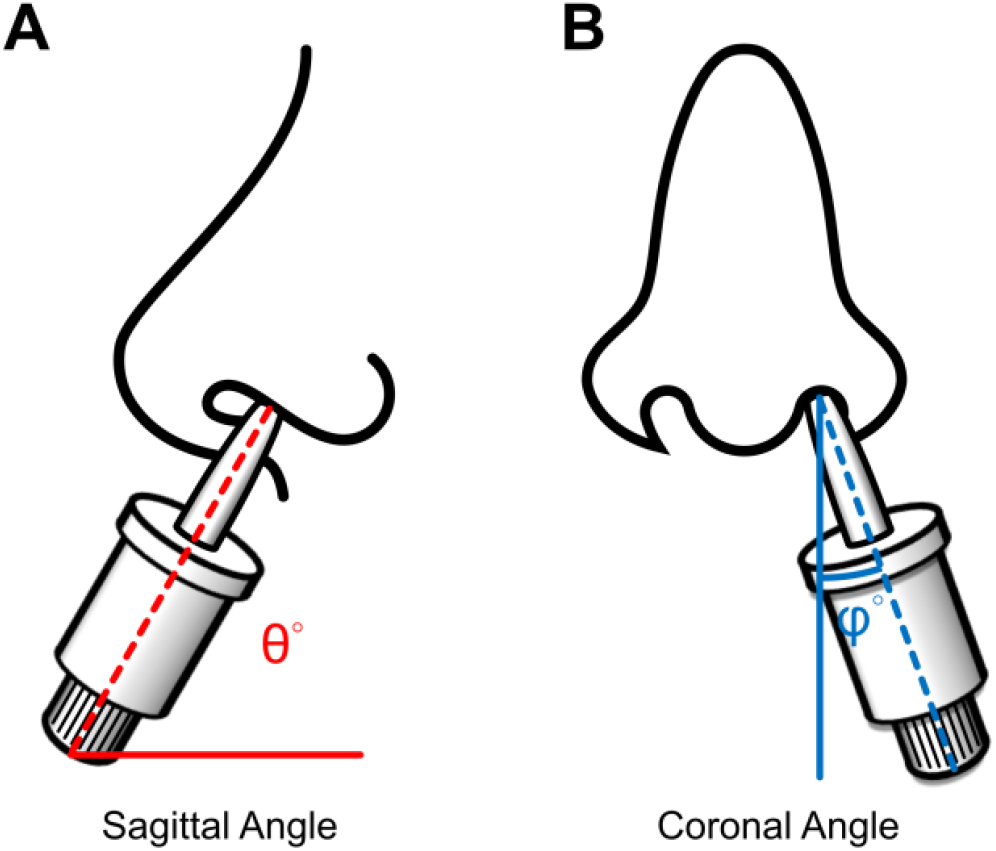
The definition of the coronal angle and the sagittal angle.

**Fig. sup2.**
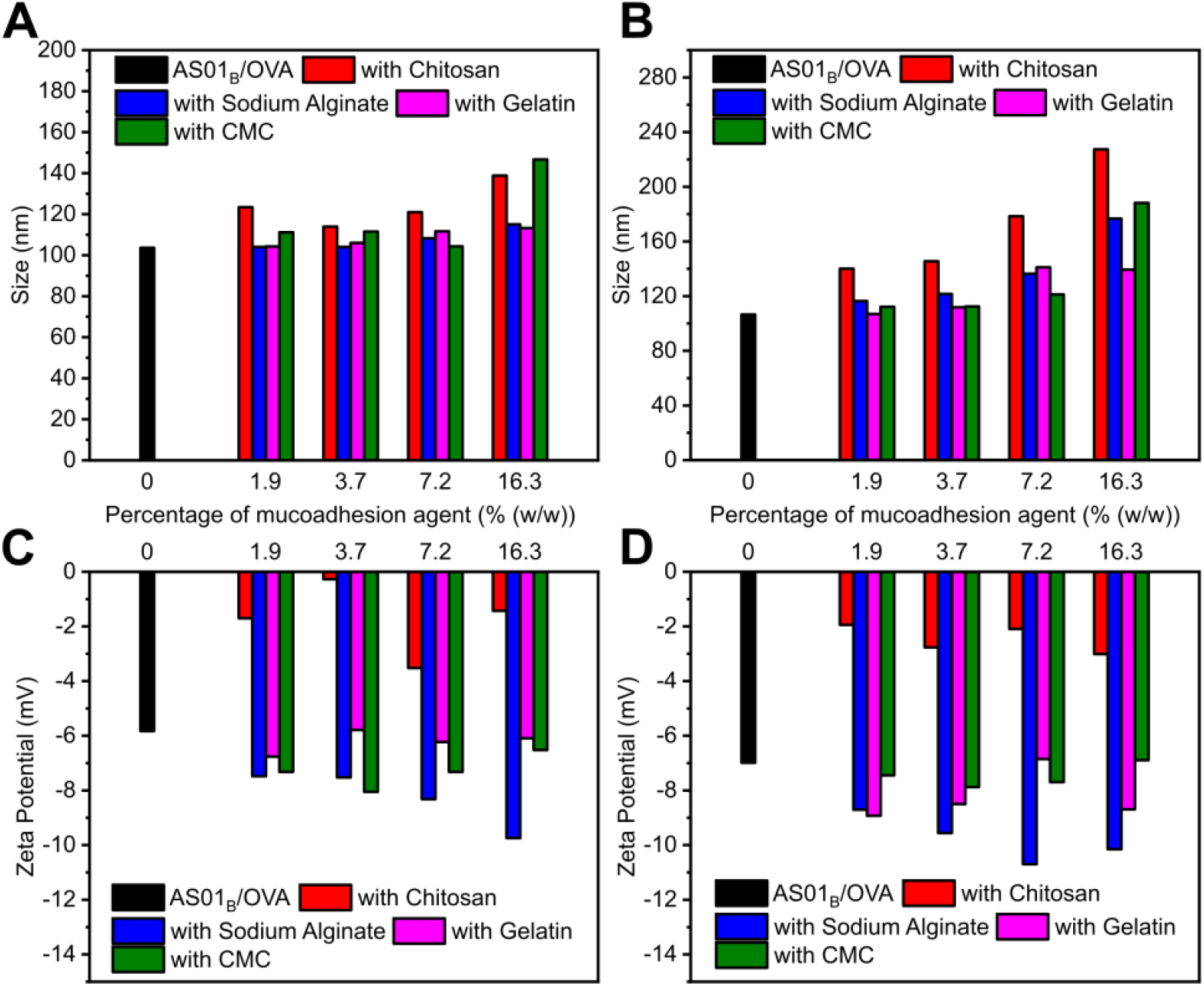
Particle size and zeta potential of AS01_B_/OVA without or with different concentrations of mucoadhesive agents before (A, C) and after being subjected to thin-film freeze-drying (B, D) (n = 1).

**Fig. sup3.**
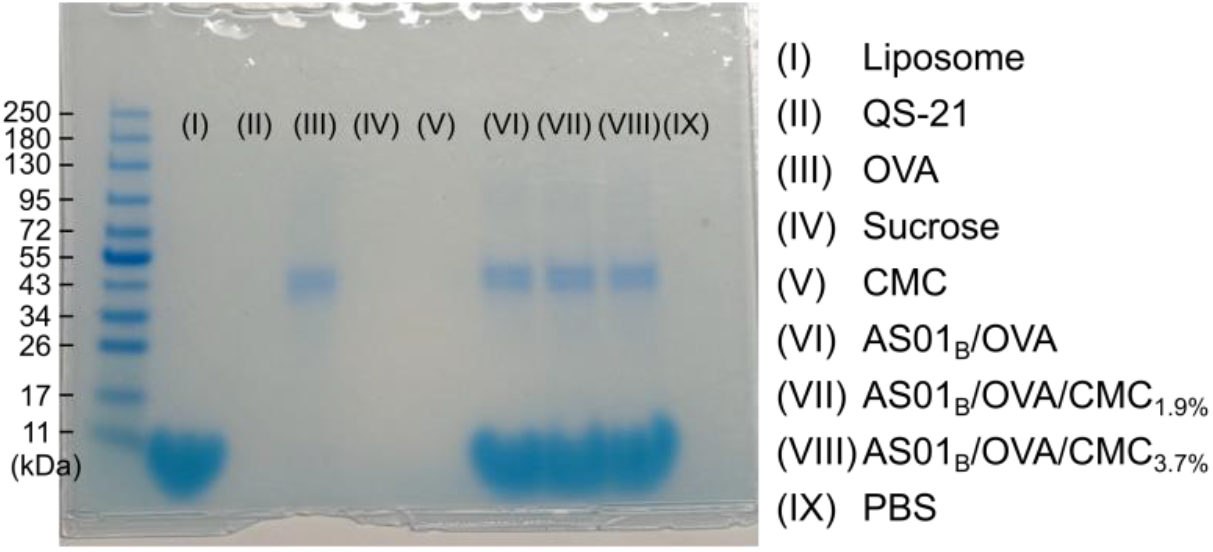
SDS-PAGE images of the AS01_B_/OVA vaccine with 0, 1.9, and 3.7% of CMC and their individual component (i.e., liposome, QS21, OVA, sucrose, CMC, PBS).

**Fig. sup4.**
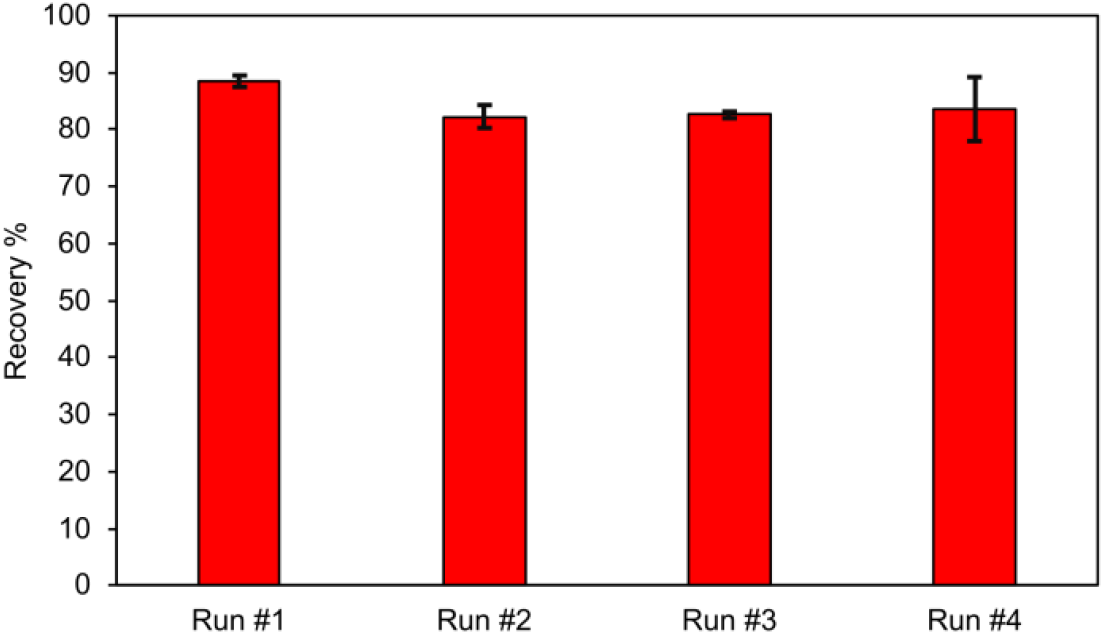
The recovery percentage of the TFF AS01_B_/OVA/CMC_1.9%_ vaccine powder in the 48-year-old male nasal cast.

**Fig. sup5.**
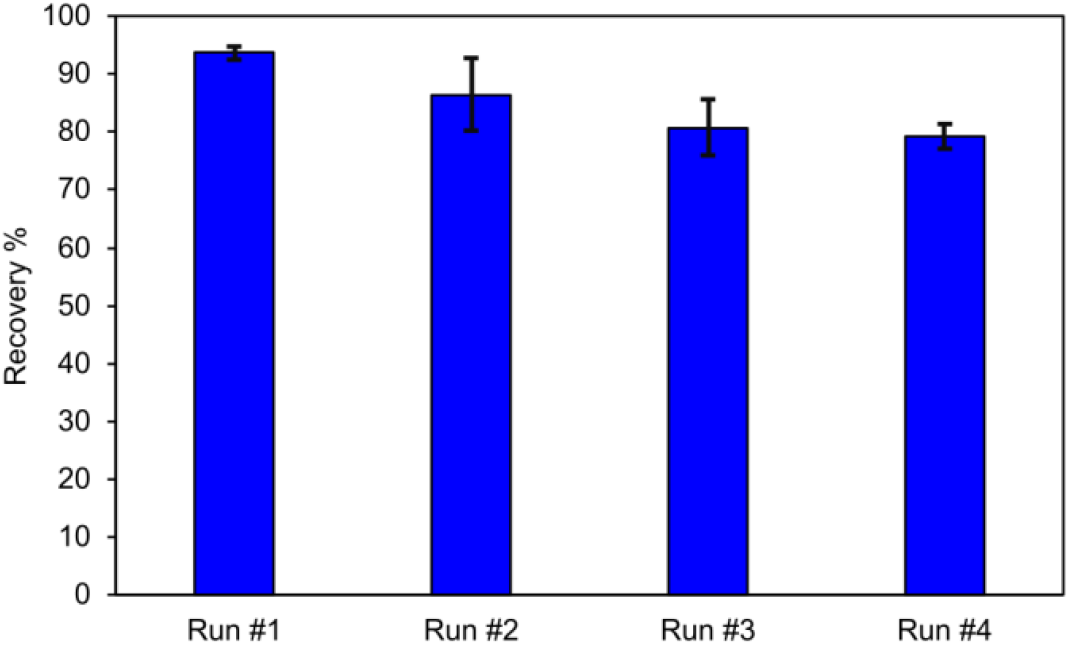
The recovery percentage of the TFF AS01_B_/OVA/CMC_1.9%_ vaccine powder in the 7-year-old female nasal cast.

**Fig. sup6.**
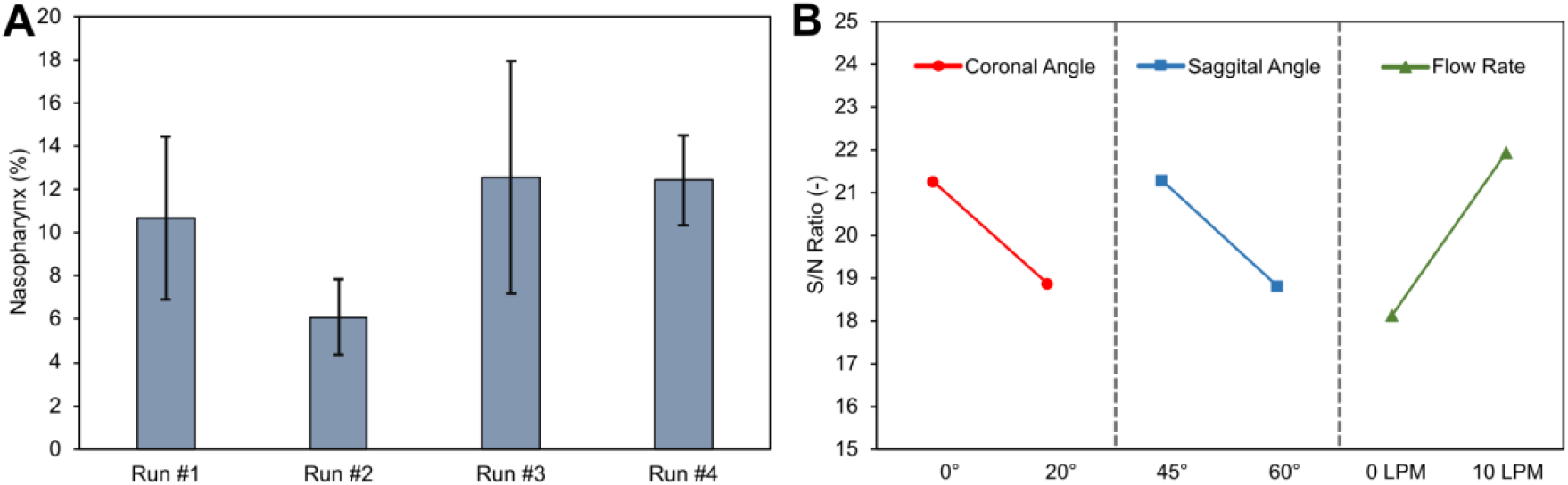
**(A)** The deposition percentage of the TFF AS01_B_/OVA/CMC_1.9%_vaccine powder in the nasopharynx region of a 48-year-old male nasal cast. (**B**) The S/N ratio of each parameter at different levels.

**Fig. sup7.**
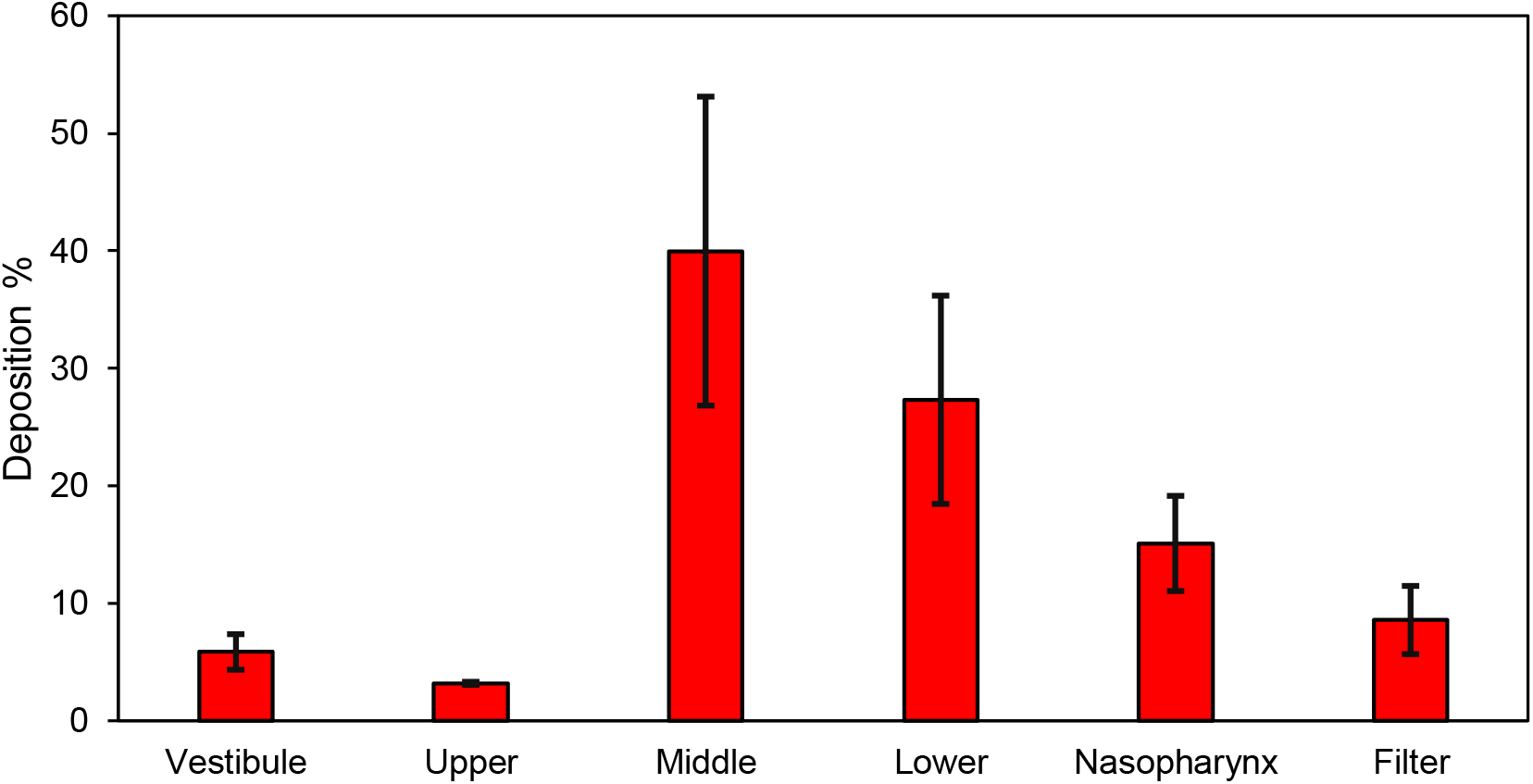
The deposition percentage of the TFF AS01_B_/OVA/CMC_1.9%_ vaccine powder in the nasopharynx region of a 48-year-old male nasal cast when applied using a UDSP nasal device at the following condition: coronal angle = 0°, sagittal angle = 45°, flow rate = 10 LPM.

**Fig. sup8.**
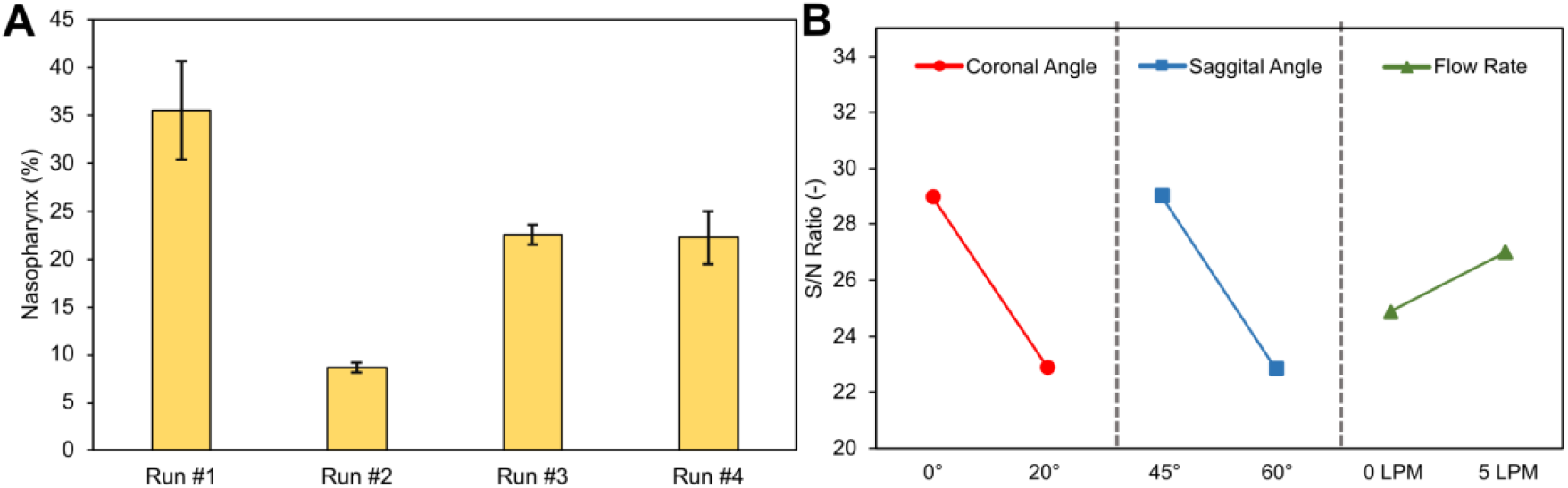
(**A**) The deposition percentage of the TFF AS01_B_/OVA/CMC_1.9%_ vaccine powder in the nasopharynx region of a 7-year-old female nasal cast. (**B**) The S/N ratio of each parameter at different levels.

